# Microglia aging in the hippocampus advances through intermediate states that drive activation and cognitive decline

**DOI:** 10.1101/2024.04.09.588665

**Authors:** Jeremy M. Shea, Saul A. Villeda

**Author notes:** Corresponding Authors: Dr. Saul Villeda, PhD University of California San Francisco 513 Parnassus Ave, Box 0452 San Francisco, CA 94143; Dr. Jeremy Shea, PhD University of California San Francisco Department of Anatomy Department of Anatomy 513 Parnassus Ave, Box 0452 San Francisco, CA 94143.

## Abstract

During aging, microglia – the resident macrophages of the brain – exhibit altered phenotypes and contribute to age-related neuroinflammation. While numerous hallmarks of age-related microglia have been elucidated, the progression from homeostasis to dysfunction during the aging process remains unresolved. To bridge this gap in knowledge, we undertook complementary cellular and molecular analyses of microglia in the mouse hippocampus across the adult lifespan and in the experimental aging model of heterochronic parabiosis. Single-cell RNA-Seq and pseudotime analysis revealed age-related transcriptional heterogeneity in hippocampal microglia and identified intermediate states of microglial aging that also emerge following heterochronic parabiosis. We tested the functionality of intermediate stress response states via TGFβ1 and translational states using pharmacological approaches *in vitro* to reveal their modulation of the progression to an activated state. Furthermore, we utilized single-cell RNA-Seq in conjunction with *in vivo* adult microglia-specific *Tgfb1* conditional genetic knockout mouse models, to demonstrate that microglia advancement through intermediate aging states drives transcriptional inflammatory activation and hippocampal-dependent cognitive decline.

## INTRODUCTION

Microglia are one of few immune cells with long term residence in the brain parenchyma^1^. Microglia support neural development by synaptic pruning and neurogenic function in the developing brain, followed by homeostatic maintenance in the adult brain^2^. Alternatively, in the aged brain microglia exhibit unbalanced phenotypes, aggravating perceived insults rather than responding productively to environmental signals^3^. While microglia have been identified as principal mediators of multiple age-related neurodegenerative pathologies^4^, mechanisms leading to microglia dysfunction during aging remain obscure.

Numerous studies have indicated that aged microglia are inflamed^5^, have reduced phagocytic capacity^6,7^, and have decreased motility^8^. Microglia exhibit several hallmarks of aging that potentially contribute to their age-related dysfunction, such as shortened telomeres^9^, altered intercellular communication^10^, molecular alterations^11^, and a loss of proteostasis^12^. Furthermore, many recent studies have started to reveal the molecular changes that define microglial aging. Single cell RNA-Seq (scRNA-Seq) analyses indicate that microglia isolated from the entire brain lose homeostasis and activate inflammatory transcriptional profiles with age^13,14^. Data rich studies have also revealed partial overlaps between aging microglia and those from disease models, including Alzheimer’s disease^15–19^. Interestingly, the microglial response to aging shows regional variation, as white matter-rich regions induce an interferon signature in aged microglia^20^. Studies using aged plasma administration and heterochronic blood exchange demonstrate that microglia aging is in part driven by the aged systemic environment ^21,22^. However, the genesis of age-related microglial phenotypes has not been extensively investigated. So, we set out to characterize the progression of age-related hippocampal microglial changes, aiming to uncover intermediate states that could be intrinsic to the aging process. To do so, we undertook complementary cellular and molecular analyses of microglia across the adult lifespan and in heterochronic parabiosis - an experimental model of aging in which the circulatory systems of young adult and aged animals are joined^23^.

In this study, we report that microglia in the adult mouse hippocampus, a brain region responsible for learning and memory and susceptible to age-related cognitive decline, advance through intermediate states that drive inflammatory activation during aging. We utilize scRNA-Seq across the adult lifespan to identify intermediate transcriptional states of microglial aging that emerge following exposure to an aged systemic environment. We observe heterogeneous spatiotemporal dynamics of microglia activation across the adult lifespan with pseudotime and immunohistochemical analysis. Functionally, we tested the role of these intermediate states using *in vitro* microglia approaches and *in vivo* temporally controlled adult microglia-specific *Tgfb1* conditional genetic knockout mouse models to demonstrate that intermediates represent modulators of the progression of microglia from homeostasis to activation, with functional implications for hippocampal-dependent cognitive decline.

## RESULTS

### Complementary single-cell transcriptional and immunohistochemical analysis reveals the dynamics of heterogeneous hippocampal microglia aging

Previously, single-cell transcriptional profiling of microglia uncovered a diversity of responses during development and in response to pathology aggregated over multiple brain regions^13,14,16,19,24,25^. Microglia have region-specific transcriptional states, so we decided to focus our analyses on the hippocampus^11^, a region with well-reported functional differences during aging^26^. Correspondingly, we investigated age-related transcriptional heterogeneity by performing scRNA-Seq on hippocampal CD11b-positive cells isolated from mice across numerous life stages at mature (6 months), middle-age (12 months), aged (18 months), and old age (24 months) using a commonly employed pooling strategy^18,27,28^ (n = 1 pool of 5 biological replicates for each age) to reveal the dynamics of microglia aging. 82% of cells isolated with CD11b were identified as microglia using canonical markers with a small subset being proliferative (Figure S1A-C). Dimensionality reduction revealed that non-proliferative microglia were grouped into interconnected clusters that contained homeostatic, transition, and activated states, along with a semi-distinct cluster with interferon activation (Figure 1a, S1D-G). Overlaying microglia ages onto the clusters revealed progression from a homeostatic state in younger microglia to an activated state in old microglia, suggesting that gradual transcriptional changes occur during microglial aging (Figure 1B,C, S1H). Furthermore, the heterogeneity of microglial transcriptional programs increased with age (Figure S1I). Expression of *Cx3cr1* and *Itgam* exemplified the homeostatic state, while *B2m*, *Apoe*, *Cd48*, and *Lyz2* depict the activated state (Figure 1D). Interestingly, several immediate early genes (*Jun*, *Klf2*, *Fos*), as well as the purinergic receptor gene *P2ry12*, exhibited transient increases in expression at middle-age (Figure 1D). Alternatively, the interferon cluster represents a small proportion of the overall microglia population with little age-related changes in the number of these cells, but augmented expression of interferon genes during aging (Figure 1A,C,D, S1G). Thus, gene expression changes exhibited unique patterns during aging, with certain genes showing age-related decreases (e.g. *Tgfbr1*) or increases (e.g. *Apoe*), while a subset of genes (e.g. *Tgfb1*) have peaks of expression at middle age (Figure 1D,E,G; Table S1). Several complement genes implicated in age-related synaptic loss^10,29^(e.g. *C1q* and *C3*) demonstrated progressive age-related transcriptional increases, and the increased complement activity in microglia was confirmed by immunohistochemistry (Figure 1E,F).

**Fig. 1.**
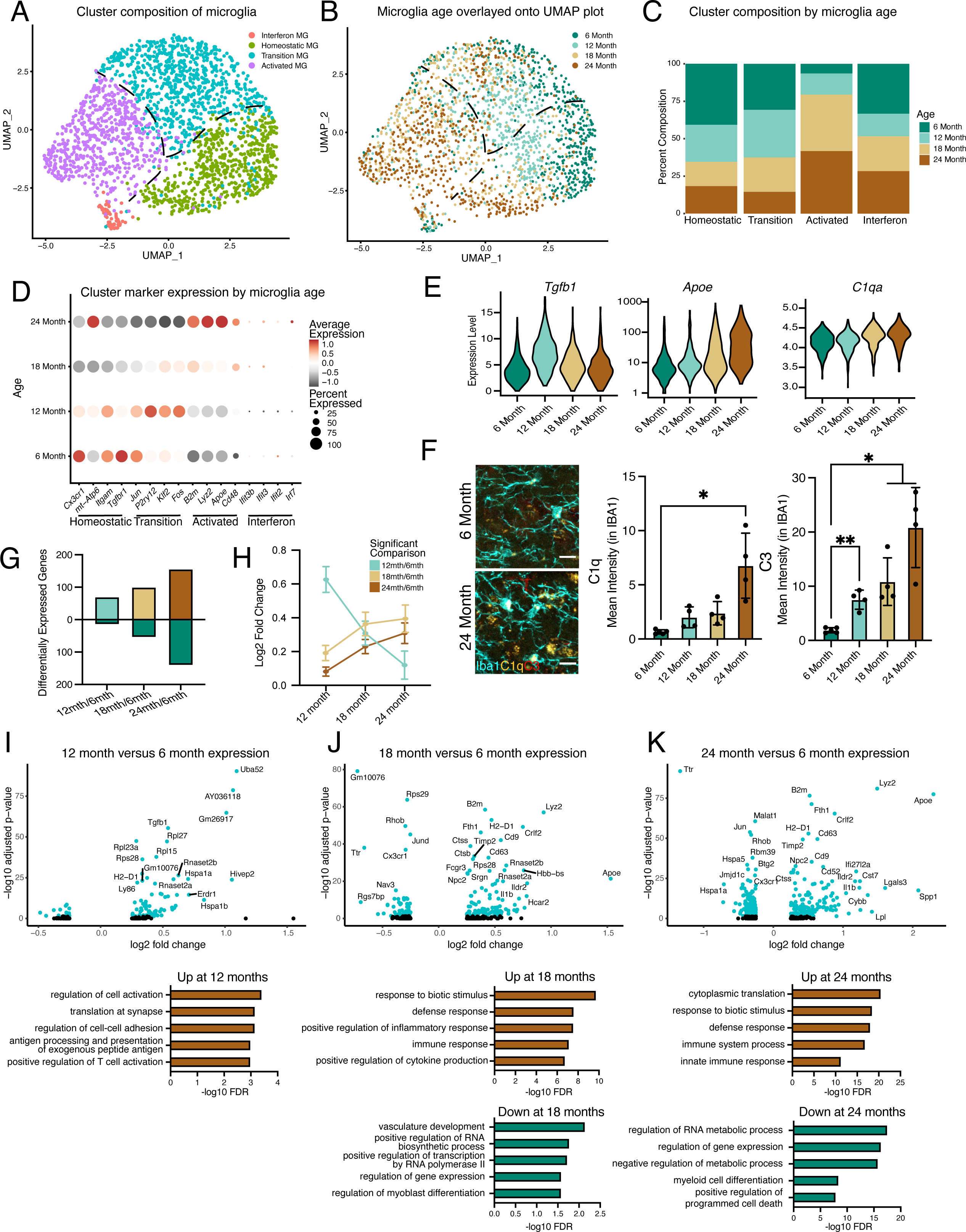
Hippocampal microglia exhibit progressive age-related transcriptional inflammatory activation. **A**, UMAP plot of microglia separated into transcriptional clusters. (n = 1 pool of 5 animals for each age) **B,** Superimposition of ages onto the UMAP plot with dashed lines signifying relative cluster demarcation. **C,** Percent composition of each cluster by age. **D,** Dot plot of expression of cluster markers sorted by age. Percent of cells expressing the gene and average normalized expression are represented. **E,** Violin plots of genes with dynamic age-related expression patterns. **F,** Representative images and quantification of C1q (yellow), C3 (red), and Iba1 (cyan) staining in the hilus and molecular layer (ML) of 6- and 24-month-old mice. (n=4-5 mice per group; mixed effects analysis followed by Dunnett’s multiple comparisons; *P<0.05, **P<0.01, ****P<0.0001). **G,** Number of differentially expressed genes from the 6-month timepoint at each age. Bars above the intersect represent increased expression and those below represent decreased expression. **H,** The average expression change at all ages for genes differentially regulated genes at individual ages represented by the color scheme in **B**. **I, J, K,** Volcano plots of differentially expressed genes in microglia between 6- and 12-months (**I**), 6- and 18-months (**J**), and 6- and 24-months (**K**) and corresponding gene ontology analysis of genes with significantly increased (brown) or decreased (green) expression for each comparison. Data are shown as mean±s.e.m.

The effects of aging on microglia could be progressive or sudden, so we investigated if we could find evidence of early aging of hippocampal microglia. We observe relatively few differentially genes between 6- and 12-month (Figure 1G,H, Table S1). The number of differentially expressed genes gradually increased through the 24-month time point (Figure 1G-J, Table S1), suggesting that hippocampal microglia progressively accumulate aged-related expression changes. Furthermore, when investigating the outcome for genes with significant changes for each age, we find that genes that are differentially expressed at 12 months lose their expression change, while genes that change at later ages show progressive expression differences (Figure 1H). GO analysis of the microglia from 6- and 12-month-old mice revealed that activation and translational programs are modestly enriched in genes with increased expression during early aging (Figure 1I). The latter stages of aging display more prominent enrichment of immune activation and translation processes in genes with increased expression, while revealing that genes that regulate transcriptional processes exhibit decreased expression (Figure 1J,K). Thus, microglia progressively age with subsets of cells at each age group being forerunners towards inflammatory activation or refractory to aging by remaining homeostatic. This analysis further suggests that microglia could pass through transitory states during aging.

As scRNA-Seq analysis revealed transcriptional heterogeneity during hippocampal microglial aging, we next assessed microglial phenotypic diversity across hippocampal subregions that could mirror the differential impact of aging across these hippocampal subregions at a functional level^30,31^. We subdivided the adult hippocampus into functional units, and quantified accumulation of puncta of the lysosomal marker CD68 in microglia to identify activation in the dentate gyrus (DG), molecular layer (ML), granule cell layer (GC), hilus, CA3, and CA1 at 3, 6, 12, 18, and 24 months of age (Figure S2A-C). We find striking spatial differences in microglial activation patterns during aging, with the GC, hilus, CA3, and outer CA1 regions showing robust microglia activation that increases beginning at middle age, while the ML and inner CA1 exhibit no age-related activation (Figure S2D-F). Additionally, we characterized the levels of the pro-inflammatory transcription factor NFKB p65 in microglia during aging^32^. Microglia-specific expression of NFKB p65 demonstrates region- and age-specific accumulation that mirrors, in part, age-related microglial CD68 increase (Figure S2G-I). These results indicate temporally defined and spatially heterogeneous microglia activation within the aging hippocampus that complements the identification of heterogeneous transcriptional activation of microglia during aging.

### Trajectory analysis indicates that aging microglia pass through intermediate states

To determine the transcriptional progression of microglia aging towards an activated state, we performed pseudotime analysis of the scRNA-Seq data using Monocle and observed several trajectories (Figure 2A). We find that the pseudotime trajectories progressed along the biological ages of the mice, indicating the suitability of this analysis for modeling aging progression in microglia (Figure 2B). We focus our analysis on one trajectory with two semi-distinct subbranches that terminated in transcriptional inflammatory activation, since the alternate branch had minimal transcriptional differences between the beginning and end of the trajectory (Figure 2B,C). To gain insight into the transcriptional states of microglial aging along this inflammatory trajectory, we used spatial autocorrelation analysis and identified five expression modules of coregulated genes (Figure 2B,C, Table S2). We named these modules according to the most enriched GO pathways and known roles of prominent genes for microglia. This trajectory proceeds from high expression of homeostatic genes and mitochondrial processes (module 1) to transient expression of stress response genes and *Tgfb1* (module 2), of which TGFβ signaling is critical for microglia development^33,34^. A concerted increase in expression of ribosomal genes (module 3) precedes one subbranch of inflammatory activation (module 4), while the other myeloid activation subbranch (module 5), characterized by *B2m* and *C1qc* expression, proceeds independently of increased ribosomal gene expression. GO analysis of genes specifically upregulated at 12-months showed an enrichment of stress response pathways (Figure S3A,B), corroborating the findings of the pseudotime analysis that intermediate stages of microglial aging pass through stress response pathways. We complemented Monocle pseudotime analysis using another distinct trajectory analysis (Scorpius) that uncovered similar progression through stress-response and translational intermediate states, followed by inflammatory activation (Figure S3C-D). We further corroborate the transient nature of the intermediate stress response module (module 2), observing KLF2 expression that peaks at 18 months of age by immunohistochemistry (Figure 2H) and *Tgfb1* expression that has a transient peak at 12 months of age by RNAscope (Figure 3B). Thus, pseudotime trajectory analysis identifies a progression of intermediate states of microglial aging.

**Fig. 2.**
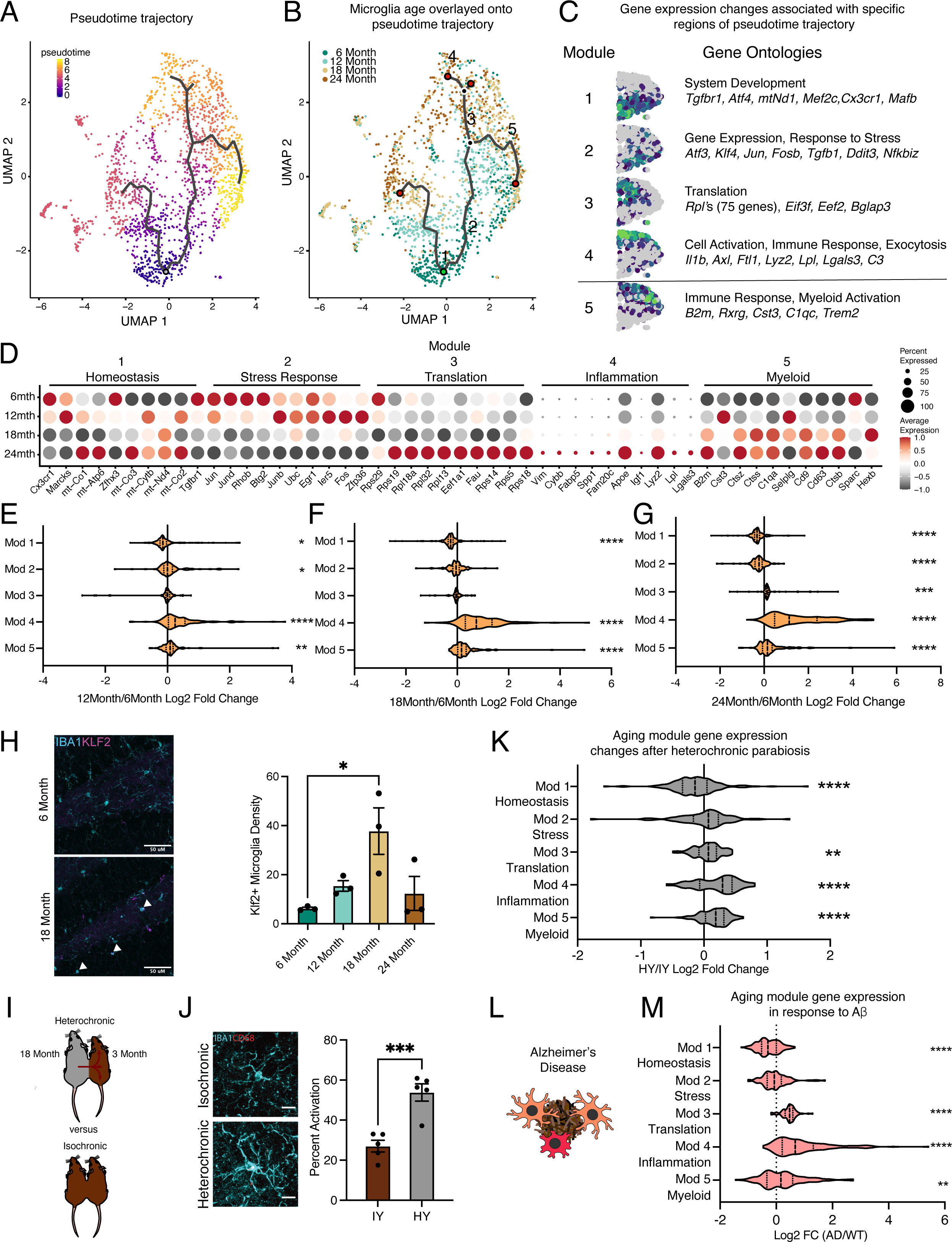
Hippocampal microglia aging advances through intermediate states that respond to systemic interventions or disease states. **A**, Pseudotime trajectories of microglia from an anchor point located in 6-month microglia presented in a UMAP plot. (n = 1 pool of 5 animals for each age) **B,** Microglia ages superimposed over pseudotime trajectories. **C,** Gene expression modules representing sections of the inflammatory aging trajectory over the right half of the UMAP plot (left). Modules were discovered using Moran’s I autocorrelation test. Top Gene Ontology terms and representative genes in each module (right). **D,** Dotplot of pseudotime modules sorted by age. Percent of cells expressing the gene and average normalized expression are represented. **E, F, G,** Average gene expression changes for each aging module represented as log2 fold change of 12-months (**E**), 18-months (**F**), or 24-months (**G**) over 6-months. **H,** Representative images and quantification of KLF2 (magenta) and IBA1 (cyan) staining in the hippocampus across ages. (n=3 mice per group; one-way ANOVA with Tukey’s post-hoc test; *P<0.05)**. I,** Diagram of the heterochronic parabiosis model with the comparisons made in scRNA-Seq. **J,** Representative images and quantification of CD68 (red) and IBA1 (cyan) staining in the hippocampus of isochronic young (IY) and heterochronic young (HY). (n=5 mice per group; unpaired Student’s T-test; ***P<0.001)**. K,** Average gene expression changes for each aging module represented as log2 fold change of heterochronic young (HY) over isochronic young (IY) adult parabionts. Data from Palovics *et al* ^38^. **L,** Diagram of microglia surrounding an Aα plaque. **M,** Average gene expression changes for each aging module represented as log2 fold change of the *App^NL-G-F^* genotype (AD) over wildtype (WT). Data from Sala Frigerio *et al* ^17^. (one sample T-test with the expected value of 0 (no change); *P<0.05, **P<0.01, ***P<0.001, ****P<0.0001). Data are shown as mean±s.e.m.

**Fig. 3.**
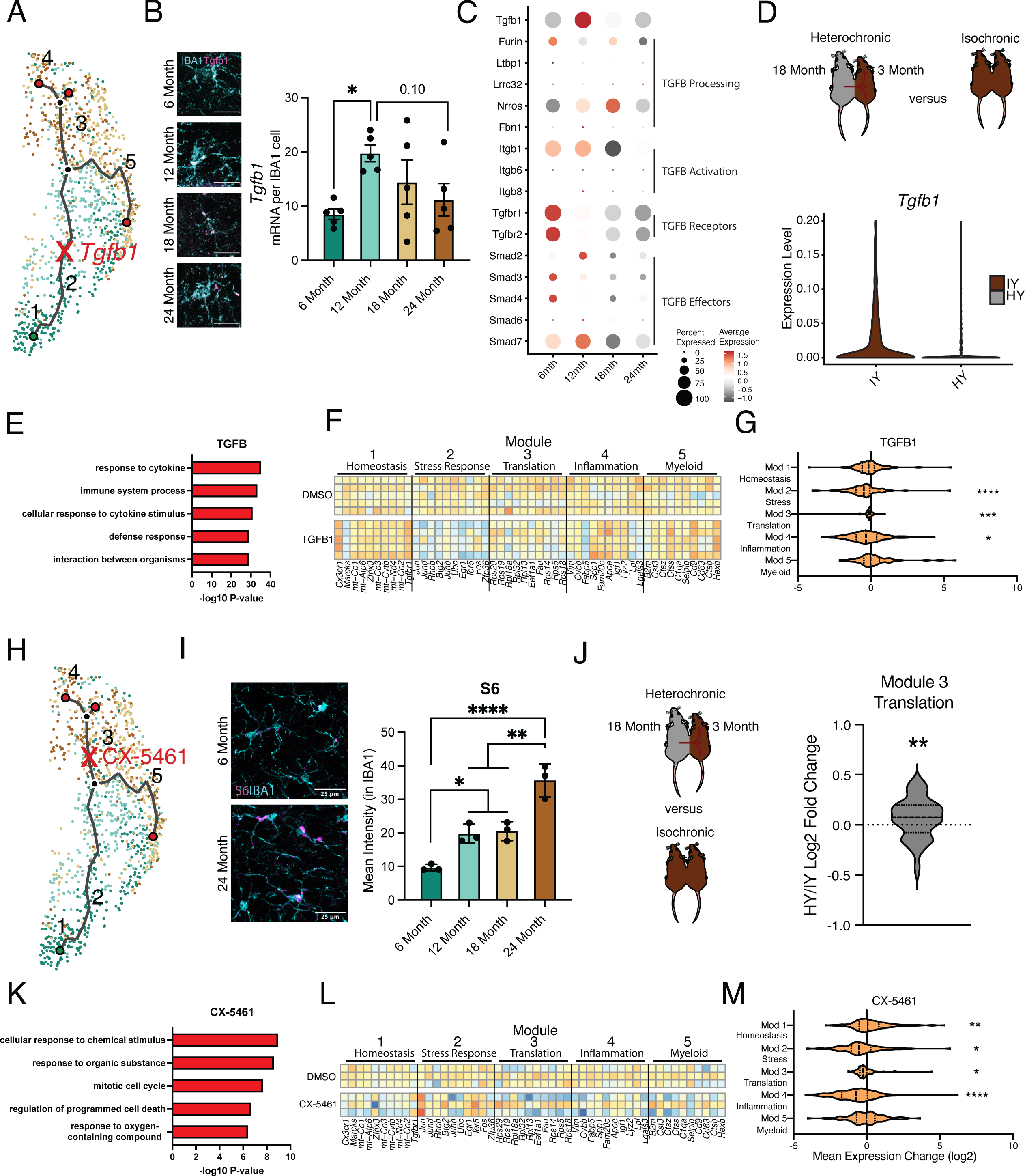
Intermediate states of microglia aging act as checkpoints on inflammatory progression. **A**, Representation of the microglia aging trajectory over the UMAP plot highlighting the region of peak *Tgfb1* expression. **B,** Representative RNAscope images and quantification of *Tgfb1* (red) expression in IBA1 (cyan) cells across ages. (n=5 per group; one-way ANOVA with Dunnett’s post-hoc test; *P<0.05)**. C,** Dotplot of the expression values of TGFB1 signaling components from scRNA-Seq of aging hippocampal microglia (6-, 12-, 18-, and 24-month-old). Percent of cells expressing the gene and average normalized expression are represented. **D,** Schematic of the heterochronic parabiosis model and quantification of hippocampal microglia expression of *Tgfb1* from isochronic young (IY) and heterochronic young (HY) adult parabionts. Data derived from Pálovics *et al*^37^. **E,** Top gene ontology terms for the set of genes with significantly decreased expression in bulk microglia RNA-Seq following TGFB1 treatment compared to control (DMSO) in LPS treated microglia. (n=5 per group) **F,** Heatmap of top 10 genes in each aging module following TGFB1 compared to DMSO in LPS treated microglia. **G,** Average gene expression changes for each aging module represented as log2 fold change of TGFB1 treatment over DMSO. (one sample T-test with the expected value of 0 (no change); *P<0.05, ***P<0.001, ****P<0.0001). **H,** Representation of the microglia aging trajectory over the UMAP plot highlighting the stage where CX-5461 modulates the trajectory. **I,** Representative images of S6 (magenta) and Iba1 (cyan) staining in the hippocampus of 6- and 24-month-old mice and quantification across aging. (n=3 mice per group; one-way ANOVA with Tukey’s post hoc test; *P<0.05, **P<0.005, ****P<0.0001). **J,** Schematic of the heterochronic parabiosis model and quantification of hippocampal microglia expression of translation module from isochronic young (IY) and heterochronic young (HY) adult parabionts. Data derived from Pálovics *et al*^37^. (one sample T-test with the expected value of 0 (no change); **P<0.01). **K,** Top gene ontology terms for the set of genes with significantly decreased expression in bulk microglia RNA-Seq following CX-5461 treatment compared to control (DMSO) in LPS treated microglia. (n=3 per group) **L,** Heatmap of top 10 genes in each aging module following CX-5461 compared to DMSO in LPS treated microglia. **M,** Average gene expression changes for each aging module represented as log2 fold change of CX-5461 treatment over DMSO. (one sample T-test with the expected value of 0 (no change); *P<0.05, **P<0.01, ****P<0.0001).

The gene level dynamics of microglia aging are further revealed when interrogating the modules at each timepoint. Expression of several genes in the homeostatic cluster, including *Cx3cr1* and *Tgfbr1*, progressively decline throughout the aging timeline (Figure 2D). Stress response gene expression peaks at early timepoints during aging, while those genes in the translation module have coherent upregulation at 24-months of age (Figure 2D). The top ranked genes in the inflammation module gradually increase their expression throughout aging in a consistent manner (Figure 2D). Alternatively, several genes in the myeloid activation module have expression peaks at 12-(*Cst3*, *Selplg*) *or* 18-months (*Ctss*, *Cd9*) of age (Figure 2D). When the modules are collapsed into metagenes, we observe that the homeostatic module is reduced early on in aging, while the stress response module follows later (Figure 2E-G). The ribosomal and inflammatory modules display augmented expression at latter ages (Figure 2E-G). Alternatively, the myeloid activation consistently has slightly increased expression throughout aging. These results suggest that with advancing age microglia lose homeostatic and stress response functions, while increasing their translational and inflammatory capacity.

Next, we used models of pro-aging systemic interventions^35^ and age-related neurodegenerative disease^36^ to determine their impact on hippocampal microglia aging trajectories. First, we used the heterochronic parabiosis model^37^ to investigate whether exposure to an aged systemic environment could promote progression of young adult microglia along the identified aging trajectory (Figure 2I). Performing immunohistochemistry for IBA1 and CD68, we reveal that an aged systemic environment activates microglia (Figure 2J). Next, we utilized publicly available scRNA-Seq datasets of parabiosis^38^ and Alzheimer’s Disease^17^ models to investigate the transcription consequences of these interventions on microglia aging trajectories. Analyzing the scRNA-Seq dataset from Palovics *et al*^38^ we find that hippocampal microglia from young adult heterochronic parabionts have decreased expression of homeostatic genes (module 1), while having increased expression of ribosomal genes (modules 3), inflammatory activation genes (module 4) and myeloid activation genes (module 5) (Figure 2K, S3E). Using scRNA-Seq from Sala Frigerio *et al*, we analyzed the *App^NL-G-F^* transgenic mouse model data at 12 months of age to determine the effects of Alzheimer’s disease pathology on microglial aging^17^ (Figure 2L). We observe exaggerated advancement along the aging trajectory in *App^NL-G-F^*mice compared to controls, as expression is shifted towards the age-associated modules (Figure 2M, S3F,G). These shifts in gene expression posit that the aged systemic and diseased environments drive adult microglia advancement along an aging-associated transcriptional trajectory.

### Intermediate states of microglia aging act as modulators of age-related trajectory progression

Next, we sought to further corroborate microglial age-related molecular changes associated with individual intermediate states in the aging hippocampus. Specifically, we assessed changes in the intermediate stress response (module 2) and translation (module 3) modules by examining expression of *Tgfb1* (Figure 3A,B) and the ribosomal protein S6 (Figure 3G,H), respectively. The vast majority (>95%) of *Tgfb1* signal localized to microglia at every age (Figure S4A), indicating microglia are the predominant source for hippocampal TGFB1. Consistent with scRNA-Seq analysis (module 2) (Figure 1E), we observed highest *Tgfb1* expression in microglia by middle-age using RNAscope (Figure 3B). TGFBR1 activation was reduced during aging in hippocampal microglia (Figure S4B), and genes involved in the TGFβ signaling pathway exhibited age-related expression changes with several having peak expression at 12 months of age (Figure 2C, S4C). Next, we examined at a transcriptional level whether TGFα1 is likely to act in an autocrine or paracrine fashion in microglia. To do so, we interrogated our scRNA-Seq dataset, leveraging the variation in microglia *Tgfb1* expression to probe the relative activity of TGFα1. High expression of downstream TGFα signaling pathway components in microglia with high *Tgfb1* expression would point to autocrine mechanisms while, alternatively, high expression of downstream TGFα signaling pathway components in microglia with low *Tgfb1* expression would point to paracrine mechanisms. We observed highest expression of TGFα signaling pathway components and targets in microglia with the highest expression of *Tgfb1* (Figure S4E-G), suggesting an autocrine mechanism of action. Additionally, consistent with an increase in translational components found during the scRNA-Seq analysis (module 3), we observed increased expression of ribosomal protein S6 (Figure 3I)^39^ by immunohistochemistry.

To complement hippocampal aging analysis, we assessed the impact of the aging systemic milieu on both the TGFα pathway (module 2) and translational components (module 3) in the young adult hippocampus following heterochronic parabiosis. Using the dataset from Palovics *et al*^38^, we observed a decrease in microglia *Tgfb1* and TGFβ pathway expression in young adult heterochronic parabionts compared to age-matched young adult isochronic controls (Figure 3C, S4D). Additionally, expression of the translation module in microglia increased following exposure to an aged systemic environment in the heterochronic parabiosis model (Figure 3J).

To investigate the role of the intermediate stress response (module 2) and translation (module 3) modules in mediating advancement through microglial activation states, we used an *in vitro* approach. Primary microglia were treated with LPS, which induced gene expression changes with a significant overlap to aging, as well as expression changes in aging modules genes (Figure S4H,I; Table S3). We administered TGFB1 to activate TGFβ signaling (module 2) (Figure 3A) and CX-5461 to inhibit RNA Pol I synthesis and interfere with translation (module 3) (Figure 3H) in the context of microglial activation. TGFB1 treatment modified the transcriptional states of LPS-treated microglia (Figure S4J) and caused decreased expression of genes in GO terms related to inflammatory immune processes after LPS stimulation (Figure 3E). TGFB1 treatment reduced expression of later aging modules (modules 2, 3 and 4) and restored expression of the homeostatic gene *Cx3cr1* (Figure 3F,G), suggesting that TGFBα signaling acts as an aging modulator during microglial stress response. Targeting translation (module 3) with CX-5461 attenuated expression of genes in GO terms related to immune processes after LPS stimulation (Figure 3K; Table S3), as well as decreased expression of genes in modules 2, 3, and 4, while restoring expression of homeostatic genes in module 1 (Figure 3L,M). Interestingly, CX-5461 treatment had no effect on genes in module 5, which is located on an independent myeloid activation trajectory from altered translational gene expression (Figure 3L,M). These *in vitro* perturbation data identify active roles for intermediate states in mediating advancement along aging-associated inflammatory trajectories.

### Mimicking age-related changes in microglia-derived TGFB1 promotes microglial advancement along an aging-associated inflammatory trajectory *in vivo*

Having uncovered roles for intermediate states in advancing microglia along aging-associated trajectories *in vitro*, we next investigated the role of individual aging modules in adult microglia *in vivo*. We elected to disrupt the transition from homeostasis to inflammatory activation by targeting microglia-derived TGFB1, a key marker of the stress response intermediate state (module 2). While TGFB1 is critical for microglia development^34,40^, the role of microglia-derived TGFB1 in regulating microglia homeostasis during aging remains to be defined.

We generated *Tgfb1^flox/flox^*, *Tgfb1^flox/wt^*, and *Tgfb1^wt/wt^* mice carrying an inducible *Cx3cr1-Cre-ER* gene, in which *Tgfb1* is excised specifically in mature (7-8 months) microglia upon tamoxifen administration (*Tgfb1* cKO, *Tgfb1* Het, and WT, respectively) (Figure 4A). To disrupt the age-related increase in microglia *Tgfb1* observed between mature and middle-age (Figure 1E and Figure 3B), mature mice were administered tamoxifen and molecular changes were assessed two months later (Figure 4A). We find that microglia activation, as measured by CD68/IBA1 immunohistochemistry, exhibited genotype-dependent effects (Figure S5A). We performed scRNA-Seq on microglia isolated from WT, *Tgfb1* Het and *Tgfb1* cKO mature mice (n = 2 pools of 3 animals per genotype), and detected genotype-dependent changes in *Tgfb1*expression levels, as well as effectors of TGFB1 signaling (Figure S5B). Dimensionality reduction and clustering of cells from scRNA-Seq revealed readily identifiable clusters based on marker expression (Figure 4B). Composition of these clusters were dependent on the microglia genotype, with WT microglia representing the largest fraction of the homeostatic cluster and *Tgfb1* cKO microglia representing the largest fraction of the activated cluster (Figure 4C, S5C). Next, we assessed whether deletion of *Tgfb1* in mature adult microglia impacted advancement along aging-

**Fig. 4.**
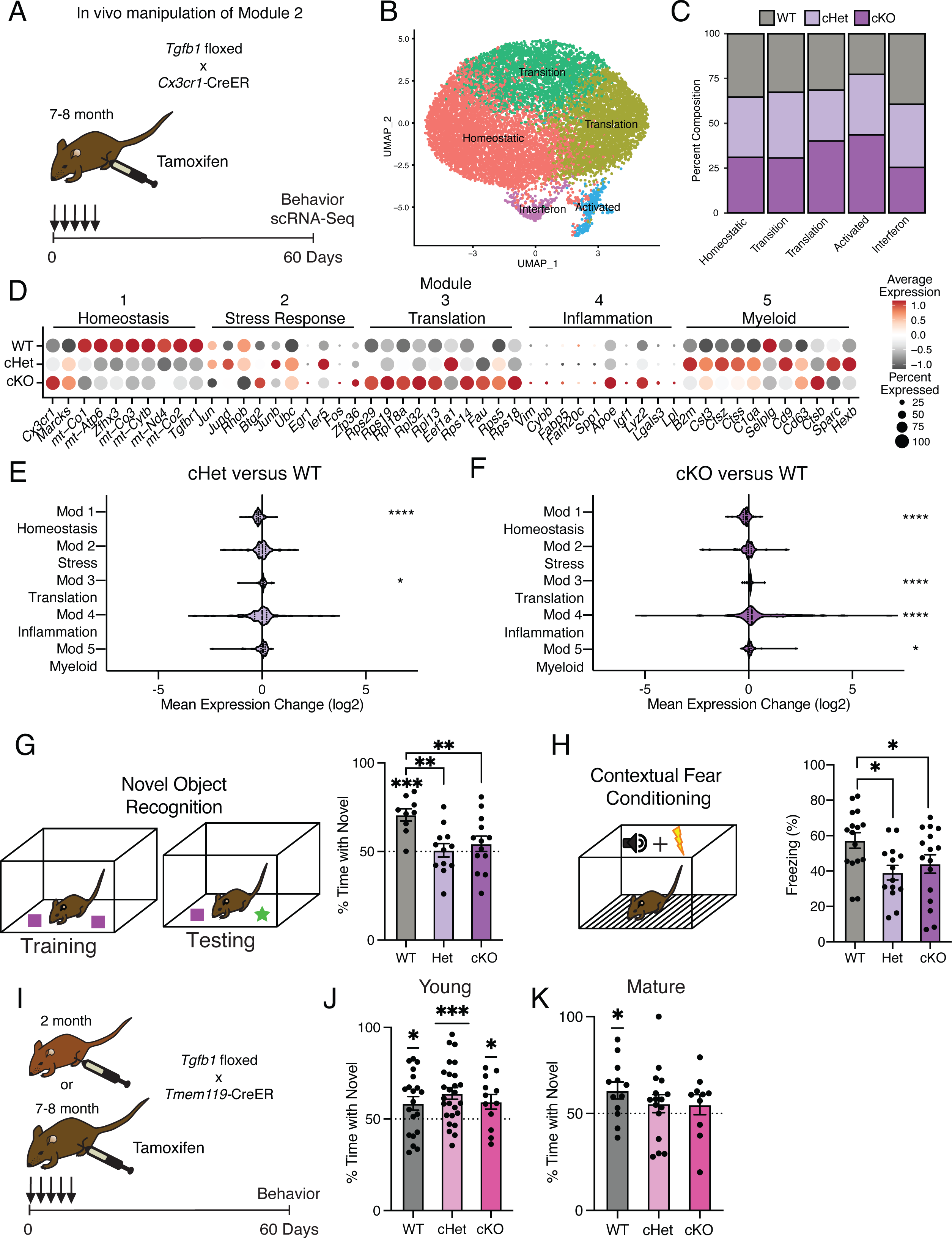
Targeting age-related changes in microglia-derived TGFB1 promotes microglia advancement along inflammatory trajectories in the hippocampus and impairs cognition. **A**, Schematic of *in vivo* manipulation schema of Module 2. Mature (7-8 months) littermate *Cx3cr1-Cre-ER^+/wt^*; *Tgfb1^flox/flox^*(cKO), *Cx3cr1-Cre-ER^+/wt^*; *Tgfb1^flow/wt^* (Het), and *Cx3cr1-Cre-ER^+/wt^*; *Tgfb1^wt/wt^* (WT) mice were administered tamoxifen and subject to hippocampal microglia scRNA-Seq and behavior analysis two months later. **B,** UMAP plot of microglia separated into transcriptional clusters. (n = 2 pools of 3 animals per genotype) **C,** Stacked bar plot of the normalized relative percentage of cells of each genotype in the identified clusters. **D,** Dot plot of expression of aging module markers sorted by genotype. Percent of cells expressing the gene and average normalized expression are represented. **E,F,** Average gene expression changes for each aging module represented as log2 fold change of either Het over WT (**E**) or KO over WT (**F**). (one sample T-test with the expected value of 0 (no change); *P<0.05, ****P<0.0001)**. G,** Novel object recognition task. (Top) Diagram of the training and testing phases of the novel object recognition paradigm. (Bottom) Quantification of the NOR testing phase represented as percentage of time spent with the novel object (over the total time spent interacting with the objects). (n=9-13 per genotype; one sample T-test with the expected value of 50 (equal time spent with each object); ***P<0.001)(differences between groups determined by one-way ANOVA; **P<0.01)**. H,** Contextual fear conditioning. (Top) Diagram of the fear conditioning paradigm. (Bottom) Quantification of the percentage of time mice froze in the contextual fear conditioning testing phase. (n=14-16 per genotype; one-way ANOVA; *P<0.05). Data are shown as mean±s.e.m. **I,** Schematic of *in vivo* manipulation of *Tgfb1* at young and mature ages. Young (2 month) or mature (7-8 months) littermate *Tmem119-Cre-ER^+/wt^*; *Tgfb1^flox/flox^* (cKO), *Tmem119-Cre-ER^+/wt^*; *Tgfb1^flow/wt^* (Het), and *Tmem119-Cre-ER^+/wt^*; *Tgfb1^wt/wt^* (WT) mice were administered tamoxifen and subject to behavioral analysis two months later. **J,K,** Quantification of the NOR testing phase represented as percentage of time spent with the novel object (over the total time spent interacting with the objects) for young (**J**) and mature (**K**) *Tmem119*-*Cre-ER*::*Tgfb1* mice. (n=12-26 in young and n=10-16 in mature mice per genotype; one sample T-test with the expected value of 50 (equal time spent with each object); *P<0.05, ***P<0.001).

associated inflammatory trajectories. Interestingly, when we analyze the effects of *Tgfb1* dosage on the aging modules, we find that the loss of a single *Tgfb1* allele reduces expression of homeostatic genes (module 1), while increasing expression of translation-related genes (module 3) (Figure 4D,E). In addition to a decrease in expression of homeostatic genes (module 1), loss of both *Tgfb1* alleles caused a greater increase in expression of translation-related genes (module 3), and led to a profound increase in inflammatory activation genes (module 4) (Figure 4D,F). Results were corroborated in an independent cohort of mature *Tgfb1* cKO mice, in which we detected altered levels of immune- and age-related cell surface markers (Figure S5D,E), transcriptional alterations that significantly overlapped with aging (Figure S5F-H; Table S4), and progression along the aging-associated inflammatory trajectories (Figure S5I,J). These data indicate that microglia-derived TGFB1 is both necessary for expression of youth-associated homeostatic genes and prevents aberrant microglia inflammatory activation, further suggesting that TGFBα signaling acts as a modulator in mediating advancement along aging-associated inflammatory trajectories.

### Mimicking age-related changes in microglia-derived TGFB1 impairs hippocampal-dependent cognitive function

It is becoming increasingly apparent that microglia interact with other cell types in the brain to influence cognitive function^2,41^. Given that manipulating the levels of microglia-derived TGFB1 altered microglia homeostasis and progression along aging-associated trajectories, we next investigated the functional consequence of the loss of microglia-derived TGFB1 on cognition. We assessed hippocampal-dependent learning and memory in WT, *Tgfb1* Het and *Tgfb1* cKO mature mice using novel object recognition and contextual fear conditioning - behavioral paradigms that are sensitive to age-related impairments^42,43^. During novel object recognition testing, WT mature mice were biased toward a novel object relative to a familiar object while neither *Tgfb1* Het or *Tgfb1* cKO mature mice showed any preference (Figure 4G). During contextual fear conditioning testing, *Tgfb1* Het and *Tgfb1* cKO mature mice exhibited less freezing compared to WT controls (Figure 4H). Alternatively, we assessed amygdala-dependent fear memory by cued fear conditioning (Figure S6A)^44^ and short-term spatial reference memory by Y maze (Figure S6B)^45^, and observed no differences across genotypes. As a control, we also profiled general health using an open field paradigm and observed no differences in total distance traveled or time spent in the center of the open field, indicative of normal motor and anxiety functions (Figure S6C).

To further examine the age-dependent role of microglia-derived TGFB1 in cognitive function, and complement our genetic approach, we utilized an alternative inducible microglia-specific *Tmem119*-CreER mouse model^46^ to manipulate *Tgfb1* expression at both young (2-3 month) and mature (7-8 month) ages (Figure 4I). We assessed hippocampal-dependent learning and memory in WT, *Tgfb1* Het and *Tgfb1* cKO young and mature mice using novel object recognition. During testing, young mice were biased toward a novel object relative to a familiar object regardless of genotype (Figure 4J), indicating normal cognitive function despite loss of microglia-specific *Tgfb1* at young age. However, while WT mature mice were biased toward a novel object relative to a familiar object, both *Tgfb1* Het or *Tgfb1* cKO mature mice lost their preference for the novel object (Figure 4K), indicating an age-dependent role for microglia-specific *Tgfb1* in maintaining cognitive function. No differences were observed in either Y maze (Figure S6D,F) or open field (Figure S6E,G) between genotypes at either young or mature ages. These behavioral data indicate that loss of microglia-derived TGFB1 not only impacts aging-associated trajectories but also promotes hippocampal-dependent cognitive impairments characteristic of age-related functional decline in an age-dependent manner.

## DISCUSSION

Cumulatively, we demonstrate that microglia aging in the hippocampus advances through intermediate states that precede inflammatory activation, with functional implications for age-related cognitive decline. These transient states, such as activation of the stress response and TGFα signaling, act as modulators of further age-related activation. Microglia demonstrate spatiotemporal dissimilarities in advancement along aging trajectories across the hippocampus. Furthermore, microglia progression through this aging trajectory is in turn influenced by the systemic environment and neurodegenerative disease states.

Analysis of our scRNA-Seq data elucidate cryptic, but necessary, intermediate states of microglia progression to age-related inflammatory activation. These states are present in middle-aged mice and constitute stress response pathways likely responsive to internal and external cellular insults^47,48^. Pseudotime analysis suggests a highly plausible sequence of molecular alterations that lead to age-related inflammatory activation of hippocampal microglia. First, mitochondrial dysfunction triggers a stress response, and products of this stress response (e.g. TGFB1) attempt to return microglia to homeostasis^49^. Here, chronic activation of the stress response leads to subsequent increased translational capacity that promotes inflammatory activation^50^. Alternatively, another branch of microglial activation advances independently of increased translational capacity, evidenced by translational inhibition having little effect on its activation following LPS stimulation. Thus, we find that increases in translational capacity predicate age-related inflammatory phenotypes, identifying potential targets to counter microglial aging. Analysis of scRNA-Seq data further identify intermediate states of microglial aging in young adult heterochronic parabionts, suggesting a pivotal role for the aged systemic environment in driving advancement towards age-related inflammatory activation.

While we find evidence of the beginnings of inflammatory activation in the hippocampus by middle age (12 months), consistent with the timing recently reported in the whole brain^14^, it should be noted that scRNA-Seq analysis of the whole brain did not observe intermediate stages in microglial aging^14^. These disparate observations could likely be the result of analyzing multiple regions with highly differentiated transcriptional programs^11^ that could obfuscate regional aging trajectories. This further underscores the need for both temporal and regional specificity in aging studies, given the level of cellular heterogeneity observed across ages and brain regions. Indeed, in the hippocampus, it is likely that the local microenvironment plays a significant role in the genesis of microglial aging, as regional distinctions in cellular function and composition – even in apposed subregions – elicit different outcomes in microglia with age and following exposure to an aged systemic environment. With age, multiple hippocampal subregions accumulate activated microglia beginning at 12 months of age, signifying that age-related alterations in microglia are initially present at middle age. Studying microglia, or other cell types, at these early stages of aging in a region-specific manner could represent useful models to investigate the ontogeny of aging without the confounding effects of accumulated functional deficits. Alternatively, the lack of age-related accumulation of activated microglia in the molecular layer and inner CA1, hippocampal subregions rich in synaptic terminals, points towards microenvironmental influences that may prove refractory to aging.

Of note, we find that an inflammatory trajectory of microglia aging can be mitigated *in vitro* by environmental TGFβ signaling, while loss of microglia-derived TGFB1 promoted progression along aging-associated inflammatory trajectories *in vivo*. Together, these data indicate that TGFα signaling is necessary for maintaining youth-associated homeostasis and represents a modulator in mediating advancement along aging-associated inflammatory trajectories. Furthermore, our results, combined with those of Bedolla *et al*^34^, strongly suggest that TGFα signals in an autocrine fashion to return microglia to a homeostatic state after encountering stressful stimuli. Functionally, loss of microglia-derived TGFB1 resulted in hippocampal-dependent cognitive impairments in an age-dependent manner, suggesting that advancement through aging-associated trajectories may facilitate age-related cognitive decline. Given that a transient increase in *Tgfb1* expression is observed in middle-aged microglia, it is possible that maintaining stress response-mediated signaling into older ages may also provide a means to delay age-related microglial dysfunction and maintain cognitive function. Alternatively, TGFβ signaling impairs microglial responses to Alzheimer’s disease pathology^51^, suggesting that the dynamic regulation of TGFβ signaling in response to aging-associated pathologies is necessary for tuning microglial responses.

While microglia have been demonstrated to mediate the establishment of cognitive function during development^2^ and play prominent roles in neurodegeneration^52^, little is known about how age-related changes in microglia affect cognitive function. The specificity of the cognitive deficits that we observe in *Tgfb1* Het and *Tgfb1* cKO mice suggests that age-related changes in microglia-derived TGFB1 predominantly impacts hippocampal-dependent cognitive decline. Furthermore, the deficits observed in *Tgfb1* heterozygous mice point to the importance of maintaining youthful levels of microglia-derived TGFB1 to counteract stressors that accrue during aging.

Altogether, our results highlight the need to investigate aging across the lifespan in a region-specific manner to reveal necessary cellular intermediates in the aging process. While the present study focuses on age-related changes in microglia, our findings have broad implications for other studies in determining the extent to which aging sequelae are conserved across different cell types in the hippocampus. Furthermore, therapeutic interventions targeting intermediate states in the aging process hold far reaching potential to counteract the development of age-related pathologies, including those promoting cognitive dysfunction.

## ACKNOWLEDGMENTS

We thank Dr. Erik Ullian for critically reading manuscript. **Funding**: This work was funded by National Institute on Aging (AG055292 (J.M.S.), AG055797 (S.A.V.), AG077816 (S.A.V.)), Simons Foundation (S.A.V). The authors declare no competing financial interests.

## AUTHOR CONTRIBUTIONS

J.M.S and S.A.V. developed concept and designed experiments. J.M.S conducted experiments, collected data, and performed analysis. J.M.S and S.A.V wrote manuscript. S.A.V supervised all aspects of the project. All authors had the opportunity to discuss results and comment on manuscript.

## METHODS

### Mice

All animal procedures were performed in accordance with protocols approved by the UCSF IACUC. Animals were housed in SPF barrier facilities and provided continuous food and water along with environmental enrichment. Aging characterizations were performed on an inhouse C57BL/6J mouse colony where 2-month-old mice were purchased from Jackson Laboratories (Stock 000664) and aged in the UCSF Parnassus barrier facility. Timed pregnant females were obtained for isolation of primary microglia from newborn pups. Male mice were used in all experiments. B6.129P2(Cg)-*Cx3cr1*^tm2.1^(cre/ERT2)^Litt^/WganJ (*Cx3cr1*Cre^ER^) (Stock 021160) and C57BL/6J-Tgfb1^em2Lutzy^/Mmjax (*Tgfb1*-flox) (Stock 65809-JAX) mice were obtained from Jackson Laboratories. Mice were bred to obtain *Cx3cr1*Cre^ER^ +/wt with either *Tgfb1* wt/wt, *Tgfb1* fl/fl. As *Cx3cr1*Cre^ER^ is a knock-in/knock-out allele for the microglia homeostatic gene *Cx3cr1*, all mice tested were *Cx3cr1*Cre^ER^ +/wt. To induce recombination and deletion of *Tgfb1* by Cre^ER^, we injected tamoxifen for 5 days at 90mg/kg daily, and mice were analyzed after 60 days.

### Parabiosis

Parabiosis surgery followed previously described procedures^53^. Mirror-image skin incisions at the left and right flanks were made through the skin, and shorter incisions were made through the peritoneum. The peritoneal openings of the adjacent parabionts were sutured together with chromic gut suture (MYCO Medical, GC635-BRC). Apposing elbow and knee joints from each parabiont were sutured together (Coated VICRYL Suture, Ethicon, J386) and the skin of each mouse was stapled (9-mm Autoclip, Clay Adams, 427631) to the skin of the adjacent parabiont. Each mouse was injected subcutaneously with Carpofen, Enrofloxacin, and Buprenex as directed for pain and monitored during recovery. For overall health and maintenance behavior, several recovery characteristics were analyzed at various times after surgery, including pair weights and grooming behavior.

### Bulk Microglia RNA-Seq

Microglia were isolated from C57BL6/J, *Tgfb1* cHet, and *Tgfb1* cKO using FACS. Briefly, mice were sedated with ketamine followed by perfusion with 30mL of ice-cold PBS. The entire brain was removed, then the hippocampus was sub-dissected. Single-cell suspensions were generated by enzyme-mediated (papain) and mechanical dissociation using Miltenyi Neural Dissociation Kits (P) (Miltenyi, 130-092-628) according to the manufacturer’s instructions. Myelin was depleted from the suspensions using Myelin Removal Beads (Miltenyi, 130-096-731). Cells were labeled with CD11b-APC (eBioscience, 17-0112-82, RRID:AB_469343) and CD45-PE (eBioscience, 12-0451-82, RRID:AB_465668), then sorted at 4°C into Tri Reagent with a FACS AriaII (BD). The gating strategy for microglia isolation is presented in Figure S1D, showing the collection of Cd11b^+^Cd45^Intermediate^ cells. During the collection of control and *Tgfb1* microglia, CD48 was detected in Cd11b^+^Cd45^Intermediate^ cells using CD48-Pacific Blue (BioLegend, 103417, RRID:AB_756139). FlowJo was used to analyze flow cytometry data. RNA was isolated from microglia using Tri Reagent (Sigma-Aldrich, T9424). RNA pellets were resuspended in 5uL TE buffer.

RNA was transformed into RNA-Seq libraries using an adapted version of the Smart-Seq2 protocol^54^. Briefly, 5ng of total RNA was reverse transcribed with SuperScript II (Thermo Fisher, 18064014) supplemented with betaine and MgCl2 using an oligo-dT RT primer (AAGCAGTGGTATCAACGCAGAGTACT(30)VN) with a PCR binding site and a template switching oligonucleotide (5′ AAGCAGTGGTATCAACGCAGAGTACATrGrG+G) with a homotypic PCR binding site. Betaine and MgCl2 were added to enhance the reaction. After reverse transcription, whole transcriptome amplification by PCR was performed for 12 cycles using KAPA HiFi HotStart ReadyMix (Roche, 7958935001) with a PCR primer (AAGCAGTGGTATCAACGCAGAGT). The PCR reaction was cleaned up with Ampure XP beads (Beckman Coulter, A63881). In addition, after the whole transcriptome PCR amplification step, qPCR was performed to determine the presence of microglia-specific transcripts (*Cx3cr1*) and housekeeping genes (*Gapdh*). The amplified DNA was diluted to a concentration of 0.5ng/ul, and subjected to tagmentation with the Illumina Nextera XT kit (Illumina, FC-131-1096). Each sample was PCR amplified with a unique set of Nextera indices. Bulk microglia RNA-Seq libraries were sequenced on an Illumina HiSeq, while *in vitro* microglia RNA-Seq libraries were sequenced on an Illumina Nova-Seq.

### RNA-Seq Analysis

FASTQ reads were pseudo aligned to the mouse transcriptome (GRCm39 cDNA from Ensembl) using kallisto^55^ with default parameters. Next, transcript abundance estimates were imported into DeSeq2^56^. Differential expression analysis was performed using DeSeq2 using the Wald significance test. Gene ontology analysis was performed with Panther^57–59^. Principal component analysis plots were generated with the plotPCA function included with DeSeq2 and graphed with ggplot2. Heatmaps were generated with the pheatmap package in R.

The overlap between gene expression differences in aging and LPS treatment were determined using genes significantly changed between the 6- and 24-month-old timepoints in the scRNA-Seq data and the control and LPS treated primary microglia. χ^2^ test was used to determine the significance of the overlap between these two datasets. The Venn diagram was generated using the VennDiagram package in R.

### 10x Genomics Single Cell RNA-Sequencing

10x Single-Cell RNA-Seq libraries were generated from Cd11b+ cells isolated from the hippocampi of an aging cohort of mice consisting of 6-, 12-, 18-, and 24-month-old C57BL/6J mice that were all collected and processed on the same day in an interspersed order. Mice were sedated with ketamine followed by perfusion with ice-cold PBS. Subsequently, the entire brain was removed from the mouse followed by sub-dissection of the hippocampus. For each age, hippocampi from five mice were pooled during the dissection step in HBSS at 4°C. Single-cell suspensions were generated by enzyme-mediated (papain) and mechanical dissociation using Miltenyi Neural Dissociation Kits (P). The papain dissociation was done at 37°C for 10 minutes with 3 trituration steps. All other processing steps were performed at 4°C. Myelin was depleted from the suspensions using Myelin Removal Beads (Miltenyi). Microglia were enriched using Cd11b magnetic beads (Miltenyi, 130-126-725). Cell viability was determined to be over 95%. Single-cell RNA-Seq 3’ libraries were generated from the cell suspension using 10x Genomics Chromium Single-Cell 3’ Solution. Libraries were sequenced on the Illumina Nova-Seq. Cell Ranger demultiplexed and mapped (using bcl2fastq) reads, followed by alignment (with STAR), and generation of single-cell expression matrices^60^.

The isolation protocol for the *Tgfb1* genetic mouse model was modified. We added transcriptional and translational inhibitors (Actinomycin D (Sigma-Aldrich, A1410), Anisomycin (Sigma-Aldrich, A9789), and Triptolide (Sigma-Aldrich, T3652) to prevent transcriptional changes due to *ex vivo* activation ^61^. We utilized the concentrations presented in Marsh *et al*. 2022^61^.

### Single-cell RNA-Seq Analysis

Seurat: The single-cell gene expression matrices generated by Cell Ranger were loaded into Seurat^62^.

Aging microglia dataset. Cells were filtered according to the following parameters: Genes > 1000 & RNA count > 10000 & mitochondrial percentage < 5. The count data was log normalized. The 2,500 most variable genes were identified using the “vst” method. The expression data was scaled and centered. Principal component analysis was performed, and the first 25 PCAs were used to identify nearest neighbors, and interconnected clusters were identified with a resolution of 0.4. UMAP was used to visualize the results. Preliminary dimensionality reduction and visualization with UMAP using 25 PCAs and 25 nearest neighbors in Seurat identified a large microglia cluster, plus sparse clusters consisting of astrocytes, vascular cells, neutrophils, and macrophages. *Hexb*, *P2ry12*, *Tmem119*, and *Slc2a5* are used as markers to identify microglia in the preliminary clustering, while *Nrxn1* (neurons), *Tm4sf1* (vasculature), *Dclk*1 (astrocytes), *Sdpr* (vasculature smooth muscle), *Abcb1a* (pericyte/vasculature), and *Foxq1* (macrophages) identified other cell types. The data was then subsetted on the non-proliferating microglial clusters (to avoid confounds associated with the cell cycle). The top 1,500 most variable genes in the subsetted microglia were identified using the “vst” method. The data was scaled and centered. Principal component analysis was performed, and the first 8 PCAs were used to identify nearest neighbors, and interconnected clusters were identified with a resolution of 0.25. UMAP was used to visualize the results using 8 PCAs and 100 nearest neighbors. Gene enrichment and differential expression analysis was performed on the filtered dataset to determine differences between ages.

*Tgfb1* cKO microglia dataset. Cells were prefiltered according to the following parameters: Hexb > 4 & Tmem119 > 1 & Cx3cr1 > 4 & Top2a < 1 & Mki67 < 1 nFeature_RNA > 1000 & nFeature_RNA < 6000 & nCount_RNA > 2000 & nCount_RNA < 40000 & percent.mt <10. Data was integrated using IntegrateData and scaled. Principal component analysis was performed, and the first 50 PCAs were used to identify nearest neighbors, and interconnected clusters were identified with a resolution of 0.35. UMAP was used to visualize the results. Preliminary dimensionality reduction and visualization with UMAP using 50 PCAs identified a large microglia cluster, plus clusters of other cell types. The non-proliferative microglia clusters were subsetted for further analysis (the macrophage cluster was differentiated from the microglia cluster based on increased expression of *Mcp1, Pf4,* and *Ms4a7*). The data was scaled and centered. Principal component analysis was performed, and the first 10 PCAs were used for UMAP dimensionality reduction and to identify nearest neighbors, and interconnected clusters were identified with a resolution of 0.20.

Monocle 3: Monocle 3 was used to generate the pseudotime analysis^63–66^. Data was imported from Seurat analysis and a Monocle dataset was generated. The dataset was preprocessed with normalization of the data and PCA generation. Next, the preprocessed data underwent further non-linear dimensionality reduction using UMAP. Cells were clustered, and trajectories were determined for cells within a cluster. For determining “pseudotime” aging of microglia, the root node of the pseudotime trajectory was manually placed in the cluster of 6-month cells. The graph segments of the pseudotime trajectories resulting in inflammatory activation were chosen for further analysis. Graph autocorrelation analysis was performed on the inflammatory trajectory using Moran’s I, which determines if genes are expressed in focal regions of graph space. A cutoff q-value of 0.005 was set as significant for focal expression. Subsequently, coregulated modules were determined using Louvain community analysis at an optimized resolution of 0.0078. The modules were then overlaid onto the UMAP plot to determine the region of the trajectory with focal expression of the module. The ten most significant genes of each module, as determined by q-value were used to construct the heatmaps in Figures 3F and 3I. The significant genes of each module were used to determine the effects of manipulations in the presence of LPS in Figure 3J.

Scorpius: Scorpius was used to generate a secondary pseudotime analysis that confirmed the results of the Monocle 3 analysis^67^. Data was imported from Seurat analysis as a SingleCellExperiment and dimensionality was reduced. A trajectory plot was generated followed by a trajectory inference. Candidate marker genes were identified using the gene_importances command. Subsequently, gene expression modules were generated with the extract_modules command and visualized using a heatmap.

### Immunohistochemistry

Mice were sedated with ketamine followed by perfusion with 30mL of ice-cold PBS. Brains were collected from PBS perfused mice and fixed overnight at 4°C in 4% paraformaldehyde in PBS. The brains were washed with PBS and immersed in 30% sucrose (in PBS), and then stored at 4°C until the brains sank. 40μM coronal sections were sliced using a microtome at -20°C. Sections were stored in cryopreservation buffer (40% PB Buffer (0.02 M Sodium Phosphate Monobasic, 0.08 M Sodium Phosphate Dibasic, pH 7.4), 30% Glycerol, 30% Ethylene Glycol) at -20°C until staining. Hippocampal sections were washed three times in TBST. Sections were permeabilized in pretreatment buffer (0.1% Triton-X in TBST) with rocking for 1 hour at room temperature. After three washes in TBST, sections were blocked in 5% goat serum for 2 hours (for IBA1/CD68 double stain) or 5% donkey serum (NFKB/Iba1 and C1q/C3/Iba1 stains). Blocking buffer was replaced with primary antibody mixture and incubated overnight at 4°C with rocking. The following primary antibodies were used: rabbit anti-IBA1 (1:1000, Wako, Cat# 019-19741, RRID:AB_839504), rat anti-CD68 (1:250, Bio-Rad, Cat# MCA1957, RRID:AB_322219), rabbit anti-NFKB p65 (1:500, Santa Cruz, Cat# sc-372, RRID:AB_632037), guinea pig anti-Iba1 (1:1000, Synaptic Systems, Cat# 234-004, RRID:AB_2493179), rabbit anti-C1q (1:1000, Abcam, Cat# ab182451, RRID:AB_2732849), rat anti-C3 (1:1000, Abcam, Cat# ab11862, RRID:AB_2066623), rabbit anti-S6 (1:500, Cell Signaling, Cat# 2217, RRID:AB_331355), and rabbit anti-KLF2 (1:250, Bioss, Cat# bs-2772R, RRID:AB_10857057). The sections were washed three times with TBST. Subsequently, secondary antibody solution was added to the sections, and incubated at room temperature with rocking for two hours. The following secondary antibodies were used: goat anti-rabbit AlexaFluor 488 (Life Technologies, Cat# A-11008, RRID:AB_143165), goat anti-rat AlexaFluor 555 (Life Technologies, Cat# A-21434, RRID:AB_2535855), donkey anti-guinea pig AlexaFluor 488 (Jackson Immuno, Cat# 706-545-148, RRID:AB_2340472), donkey anti-rabbit AlexaFluor 555 (Life Technologies, Cat# A-31572, RRID:AB_162543), and donkey anti-rat AlexaFluor 647 (Jackson Immuno, Cat# 712-606-150, RRID:AB_2340695). The sections were washed three times with TBST, with the first wash containing Hoescht 33342 (Invitrogen, H3570) at a concentration of 1μg/mL. The sections were transferred to phosphate buffer and mounted on frosted slides. Coverslips were mounted with Prolong Gold Antifade (Thermo Fisher, P10144) reagent after the sections were dried. Slides were imaged at 20x magnification on a Zeiss LSM 800 or LSM900 for a final resolution of 0.312μM/pixel. 8-9 z-planes were imaged at 3μM intervals. The hippocampus was imaged, and the tiled images were stitched together with Zeiss ZenPro software. 2-3 hippocampal sections were imaged from each sample.

Images were analyzed with FIJI. Maximum intensity projections of Z-stacked images were constructed. Images were converted to 8-bit images and consistent adjustments were made for each channel within an experiment. The Iba1 channel was used to quantify microglia using the particle counter function in FIJI and the counts were validated by manual visual counting. The presence of CD68, NFKB(p65), C3, or C1q in microglia was determined by creating ROIs within the microglia in a field using the create selection function on the Iba1 images. The ROIs of the hippocampal subregions and the microglia were combined, and CD68 puncta meeting a minimum particle size threshold (100 pixels, equating to 31.2μM) within microglia determined if the microglia were activated (CD68+IBA1+), while the measure function determined the intensity of NFKB(p65), C3, or C1q. To determine percent activation of microglia with IBA1/CD68 staining, we divided the number of microglia that were activated (CD68+IBA1+) by the total number of microglia (IBA1+) in the analyzed regions and multiplied by 100.

Additionally, we tested whether autofluorescence from aged tissue was interfering with our signal using TrueBlack® Lipofuscin Autofluorescence Quencher (Cell Signaling, Cat #92401). Using the IBA1/CD68 doublestain, we obtained very similary results for microglia activation. Furthermore, our KLF2 stain was performed with TrueBlack.

### *In vitro* treatment of microglia

Mixed glial cultures were generated from p1 male pups. Cortices and hippocampi were isolated from the pups, and meninges were removed. The tissue was broken up by repeatedly pipetting up and down, followed by dissociation with Trypsin. Single cell suspensions were plated on poly-L-lysine coated flasks in DMEM (Thermo Fisher, 11965-118) supplemented with 10% FBS (Gemini Bio-Products, 900-208) and antibiotic mix (Penicillin/Streptomycin)(Genesee Scientific, 25-512) and incubated at 37°C in 5% CO2. Media was changed after 4 days to remove debris. Microglia were isolated after 14 days by agitating the plates to dislodge microglia, then transferring the supernatant of each sample evenly amongst 4 wells of a 24 well plate. Microglia were incubated for 16 hours followed by replacement with serum free macrophage SFM media (Thermo Fisher, 12065074) supplemented with M-CSF (10ng/m) (Peprotech, 315-02) and antibiotic mix. After 24 hours microglia were treated with RNA polymerase I Inhibitor 2, CX-5461 (100nM)(EMD Millipore, 509265) or human TGFβ1 recombinant protein (10ng/mL)(ThermoFisher Scientific, PHG9204) for 24 hours followed by addition of lipopolysaccharides (LPS, 200ng/uL) from *E. coli* (Sigma-Aldrich, L2018) for 8 hours and ATP (10nM)(InvivoGen, tlrl-atpl) for 30 minutes. Tri Reagent was added to the culture plate, and the plates were shaken for 5 minutes. RNA was isolated according to manufacturer’s instructions. RNA-Seq libraries were generated as above for the bulk RNA-Seq.

### RNAscope

RNAscope was performed with the RNAscope® Multiplex Fluorescent Reagent Kit v2 (ACD, 323110). 40μM coronal sections were washed with TBST, then incubated with hydrogen peroxide for 45 minutes at 25°C. After washing with TBST, the sections were incubated in Target Retrieval Buffer at 95°C for 10 minutes. The sections were mounted on slides and dried overnight. The sections were treated with RNAscope Protease III for 10 minutes at 40°C. Sections were washed in water four times. Sections were incubated with *Tgfb1* probe (ACD, 443571-C2) for 2 hours at 40°C. The sections were washed twice with RNAscope wash buffer. The sections were incubated with AMP1 for 30 minutes, AMP2 for 30 minutes, AMP3 for 15 minutes, HRP-C2 for 15 minutes, TSA Plus Cy3 (1:1000, Perkin Elmer, SKU NEL744001KT) for 30 minutes, and HRP-Blocker for 15 minutes; all at 40°C with washes with RNAscope wash buffer between every incubation. RNA-Protein Co-Detection Ancillary Kit (ACD, 323180) buffer was used to block the sections before incubating with rabbit anti-IBA1 (1:500, Wako, Cat# 019-19741, RRID:AB_839504) overnight at 4°C. Sections were washed three times in TBST. Sections were incubated with secondary antibody solution with goat anti-rabbit AlexaFluor 488 (1:1000, Life Technologies, Cat# A-11008, RRID:AB_143165) for two hours at room temperature. The sections were washed three times with TBST, with the first wash containing Hoescht 33342 (Invitrogen, H3570) at a concentration of 1μg/mL Coverslips were mounted with Prolong Gold Antifade (Thermo Fisher, P10144) reagent after the sections were dried. Slides were imaged at 40x magnification on a Zeiss LSM 900. 10 z-planes were imaged at 1μM intervals.

Images were analyzed with FIJI. Maximum intensity projections of Z-stacked images were constructed. Images were converted to 8-bit images and the *Tgfb1* was thresholded. For each section, the intensity of 5-7 individual *Tgfb1* puncta were averaged to get the average puncta intensity of that section. The intensity of all *Tgfb1* signal in clearly identifiable individual IBA1 cells was measured for 20-25 cells in the molecular layer of the hippocampus. The mean *Tgfb1* count per microglia for each sample was calculated by dividing the average *Tgfb1* intensity in IBA1 cells by the average puncta intensity.

### Contextual fear conditioning

In this task, mice learned to associate the environmental context (fear conditioning chamber) with an aversive stimulus (mild foot shock; unconditioned stimulus, US) during the training phase enabling testing for hippocampal-dependent contextual fear conditioning. To also assess amygdala-dependent cued fear conditioning, the mild foot shock was paired with a light and tone cue (conditioned stimulus, CS) during the training phase. Conditioned fear was displayed as freezing behavior. Specific training parameters are as follows: tone duration is 30 seconds; level is 70 dB, 2 kHz; shock duration is 2 seconds; intensity is 0.6 mA. This intensity is not painful and can easily be tolerated but will generate an unpleasant feeling. More specifically, on day 1 each mouse was placed in a fear-conditioning chamber and allowed to explore for 2 min before delivery of a 30-second tone (70 dB) and light ending with a 2-second foot shock (0.6 mA). Two minutes later, a second CS-US pair was delivered. On day 2, each mouse was first placed in the fear-conditioning chamber containing the same exact context, but with no CS or foot shock. Freezing was analyzed for 2 minutes and represented as the percentage of time that the mouse froze over the 2 minutes. One hour later, the mice were placed in a new context containing a different odor, floor texture, chamber walls and shape. Animals were allowed to explore for 2 minutes before being re-exposed to the CS. Freezing was analyzed for 30 seconds following the CS and represented as the percentage of time that the mouse froze over those 30 seconds. Determination of freezing behavior was performed using FreezeScan video tracking system and software (Cleversys, Inc). Single outliers were removed using the extreme studentized deviate method (Grubbs’ test) with an alpha of 0.05.

### Novel object recognition

The novel object recognition task was adapted from a previously described protocol^68^. Specifically, during the habituation phase (day 1), mice could freely explore an empty open field arena for 10 minutes (motor activity and anxiety (time in center) were measured during this phase). During the training phase (day 2), two identical objects were placed in the habituated arena, and mice could explore the objects for 5 minutes. For the testing phase (day 3), one object was replaced with a novel object, and mice could explore the objects for 5 minutes. Time spent exploring each object was quantified using the Smart Video Tracking Software (Panlab; Harvard Apparatus). Two different sets of objects are used. To control for any inherent object preference, half of the mice are exposed to object A as their novel object and half to object B. To control for any potential object-independent location preference, the location of the novel object relative to the trained object is also varied. The objects were chosen based on their ability to capture the animal’s interest, independent of genetic background or age. To determine percent time with novel object, we calculate (Time with novel object)/(Time with Trained Object + Time with Novel Object) * 100. In this preference index, 100% indicates full preference for the novel object, and 0% indicates full preference for the trained object. A mouse with a value of 50% would have spent equal time exploring both objects. Mice that did not explore both objects for 5 seconds during the training phase or testing phase were excluded from analysis. Single outliers were removed using the extreme studentized deviate method (Grubbs’ test) with an alpha of 0.05.

### Y maze

The Y Maze task was conducted using an established forced alternation protocol^69^. During the training phase, mice were placed in the start arm facing the wall and allowed to explore the start and trained arm for 5 minutes, while entry to the 3rd arm (novel arm) was blocked. The maze was cleaned between each mouse to remove odor cues, and the trained arm was alternated between mice. The mouse was then removed to its home cage. After 30 minutes, the block was removed, and the mouse was returned to the start arm and allowed to explore all 3 arms for 5 minutes. Time spent in each arm was quantified using the Smart Video Tracking Software (Panlab; Harvard Apparatus). The percent time in the novel arm was defined as time in the novel arm divided by time spent in the novel and trained arms during the task.

### Datasets

All RNA-Seq and scRNA-Seq data have been deposited in the Gene Expression Omnibus and are publicly available as of the date of publication. The following datasets were generated for this manuscript and deposited in Gene Expression Omnibus: aging single-cell RNA-Seq (GSE179358), *in vitro* treated primary microglia (GSE179611), and *Tgfb1* cKO microglia RNA-Seq (GSE190007).

### Statistics

Researchers were blinded throughout histological assessments with groups being un-blinded at the end of each experiment upon statistical analysis. Data are expressed as mean ± s.e.m with individual sample values being shown. Statistical analysis was performed with Prism 8.0 (GraphPad), R, DESeq2, Seurat, or Monocle 3. Comparisons of means in histology experiments were analyzed with multiple T-tests followed by Holm-Sidak correction or mixed effects analysis followed by Dunnett’s multiple comparisons (Prism). Changes in expression for aggregated gene sets were determined with one sample T-tests with the expected value of 0 in log2 space (null hypothesis of no change). Differential expression analysis in DESeq2 was based on Wald’s significance test of the negative binomial distribution. Differential expression analysis of single-cell RNA-Seq data in Seurat was performed using non-parametric Wilcoxon rank sum tests. Autocorrelation analysis to determine focal expression Monocle 3 was done using Moran’s I. The significance of gene set overlap was determined using the χ^2^ test with R. All data generated or analyzed in this study are included in this article.

## SUPPLEMENTAL MATERIAL

**Fig. S1 Hippocampal microglia exhibit age-related heterogeneity during aging. A,** UMAP plot of all Cd11b+ cells. Several adjacent clusters of microglia were identified, as well as a cluster of proliferating microglia. Smaller populations of peripheral immune cell types – macrophages and neutrophils – were identified. Clusters of astrocytes and vascular cells were also found. Overall, greater than 82% of cells were microglia. (n = 1 pool of 5 animals for each age) **B,** Dot plot showing expression (average expression and percent of cells expressing) of top two markers for each cluster. **C,** UMAP plots with expression levels of microglia markers superimposed onto cells. Notice that peripheral immune cells express microglia markers; however, they are distinguished from microglia based on marker expression from (b). **D-G,** Volcano plots of differential gene expression for the clusters identified in Figure 1A compared to every other cluster for Homeostatic (**D**), Transition (**E**), Activation (**F**), and Interferon (**G**) microglia clusters. **H,** Cluster composition of non-proliferating microglia by age. **I,** Standardized variation of non-proliferating microglia for each age.

**Fig. S2 Spatiotemporal kinetics of microglial inflammatory activation in the aging hippocampus. A,** Diagram depicting ages utilized for immunohistochemical analysis. **B,** Diagram of the hippocampus labeled with the regions analyzed. **C,** Illustration of subregions analyzed by immunohistochemistry. **D,** Representative images and quantification of IBA1 (cyan)/CD68 (red)-positive microglia across hippocampal subregions in 3- and 24-month old mice. Scale bars are 10μM. (n=5 per group; T-test with Holm-Sidak correction; *P<0.05, **P<0.01) **E,** Heatmap of the quantification of activated microglia across ages and subregions. **F,** Representative wide field images of IBA1 (cyan)/CD68 (red) in the dentate gyrus in 3- and 24-month old mice. Scale bars are 100μM. **G,** Representative wide field images of Iba1 (cyan)/ NFKB p65 (yellow) in the dentate gyrus in 3- and 24-month old mice. Scale bars are 100μM. **H,** Representative images and quantification of Iba1 (cyan)/ NFKB p65 (yellow) staining across hippocampal subregions. Scale bars are 10μM. (n=5 per group; T-test with Holm-Sidak correction; *P<0.05, **P<0.01, ****P<0.0001) **I,** Heatmap of the quantification of NFKB signal in microglia across ages and subregions. n=4-5 mice per condition.

**Figure S3 Hippocampal microglia aging advances through intermediate states that respond to systemic interventions or disease states**. **A**, Volcano plot of differential gene expression of 12-month microglia versus all other ages with significant genes in teal. **B**, Gene ontology analysis of biological processes enriched in those genes with increased expression at 12-months of age. **C**, Representative images and quantification of CD68 (red) and IBA1 (cyan) staining in the hippocampus of isochronic young (IY) and heterochronic young (HY) along with a diagram of the comparisons. (n=5 mice per group; unpaired Student’s T-test; ***P<0.001). Data are shown as mean±s.e.m. **D,** Dotplot of pseudotime modules in young (Y), isochronic young (IY), and heterochronic young (HY) parabiont microglia. Data is from Palovics *et al*^38^. Percent of cells expressing the gene and average normalized expression are represented. **E,** Volcano plots of differential gene expression of *App^NL-G-F^* genotype (AD) over wildtype (WT) microglia at 6- (left) and 12-months of age. Significant genes are in teal. Data is from Frigerio^70^. **F,** Dotplot of pseudotime modules across ages (3-, 6-, 12-, and 21-months old) and genotypes (*App^NL-G-F^* genotype (AD) over C57Bl/6 (WT)). Data is from Frigerio^17^. Percent of cells expressing the gene and average normalized expression are represented.

**Fig. S4 Intermediate states of microglia aging act as checkpoints on inflammatory progression. A,** Quantification of percentage of *Tgfb1* RNAscope signal within IBA1 cells. **B**, Quantification of IHC of pTGFBR1 signal in IBA1 cells across ages. (n=4-5 mice per group; one-way ANOVA; *P<0.05) **C,** Quantification of the expression of TGFB1 signaling components from Figure 3C. Expression values for the genes are represented as Z-scores. (n=12 genes with reads in >25% of cells; Friedman test followed by Dunn’s multiple comparisons test; *P<0.05, ***P<0.001) **D**, Dotplot of the expression values of TGFB1 signaling components in young (Y), aged (A), isochronic young (IY), and heterochronic young (HY) parabiont microglia. Data is from Palovics *et al*^38^. Percent of cells expressing the gene and average normalized expression are represented. UMAP plot of *Tgfb1* scRNA-Seq colored by genotype. **E**, Dotplot of the expression values of TGFB1 signaling components in aging microglia categorized by the expression of *Tgfb1* into quartiles with Q1 having the lowest expression of *Tgfb1* and Q4 having the highest expression. **F,** Quantification of the expression of TGFB1 signaling components from (**E**). Expression values for the genes are represented as Z-scores. (n=12 genes with reads in >25% of cells; Friedman test followed by Dunn’s multiple comparisons test; *P<0.05, ***P<0.001) **G,** Dotplot of the expression values of selected TGFB signaling targets (activated and repressed, genes found in both Butovsky *et al.*^33^ and Qin *et al*.^71^) in aging microglia categorized by the expression of *Tgfb1*. **H,** Heatmap of gene expression changes in the top 10 genes in each aging module induced by LPS. **I**, Overlap between microglia gene expression changes induced by LPS and aging. (χ^2^<2.2e-16). **J**, PCA plot of pharmacological manipulations with or without LPS treatment.

**Fig. S5 Targeting age-related changes in microglia-derived TGFB1 promotes microglia advancement along inflammatory trajectories in the hippocampus. A**, Representative images and quantification of IBA1 (cyan)/CD68 (red)-positive microglia in wildtype, *Tgfb1* cHet, and *Tgfb1* cKO hippocampi. (n=3-5; one way ANOVA with Tukey post hoc test; *P<0.05) **B,** Dotplot of the expression values of TGFB1 signaling components from scRNA-Seq of *Tgfb1* WT, cHet, and cKO microglia. (n = 2 pools of 3 animals per genotype) **C,** UMAP plot of *Tgfb1* scRNA-Seq colored by genotype. **D,** FACS plots of gating strategy for microglia isolation for RNA-Seq in cHet and cKO hippocampi. Quantification of the flow cytometry analysis of CD11b and CD45. (n=3-5; mixed effects analysis; *P<0.05, ****P<0.0001). **E**, Example histogram and quantification of CD48 in control and *Tgfb1* cKO FACS-sorted microglia. (n=3-5 per group; T-test; ****P<0.0001). **F,** Top gene ontology terms associated with genes significantly increased in microglia bulk RNA-Seq in *Tgfb1* cKO samples (n=3-5 samples per genotype). **G**, Top gene ontology terms associated with genes significantly decreased in microglia bulk RNA-Seq in *Tgfb1* cKO samples. **H**, Overlap between concordant microglia gene expression changes induced by *Tgfb1* knockout and aging. (χ^2^<2.2e-16). **I**, Heatmap of gene expression of the top 10 genes in each aging module in control and *Tgfb1* cKO microglia. **J**, Average gene expression changes for each aging module represented as log2 fold change of *Tgfb1* cKO over control. (one sample T-test with the expected value of 0 (no change); ****P<0.0001). Data are shown as means±s.e.m.

**Fig. S6 Targeting age-related changes in microglia-derived TGFB1 impairs cognition. A**, Freezing during the cued fear conditioning phase of the fear conditioning test for the *Cx3cr1-Cre-ER*::*Tgfb1* cohort. The percent freezing was calculated for the last 30 seconds of the test following the conditioned stimulus. (n=13-16 per genotype) **B**, Diagram and results of Y-maze test for the *Cx3cr1-Cre-ER*::*Tgfb1* cohort. Results are presented as preference for the novel arm versus the trained armed (time in novel arm/ (time in novel arm + trained arm) x 100). (n=14-16 per genotype) **C**, Percent time in the center of the open field and distance traveled during the open field test for the *Cx3cr1-Cre-ER*::*Tgfb1* cohort. (n=14-16 per genotype) **D, F,** Results of Y-maze test for the young (**D**) and mature (**F**) *Tmem119-Cre-ER*::*Tgfb1* cohort. (n=8-16 per genotype for young and n=10-16 per genotype for mature) **E,G,** Percent time in the center of the open field and distance traveled during the open field test for the *Tmem119-Cre-ER*::*Tgfb1* cohort. (n=12-26 per genotype for young and n=10-16 per genotype for mature) Data are shown as mean±s.e.m.

**Table S1 Gene markers for each age group and differential expression between the 6- and 12-, 18-, or 24-month-old microglia in single-cell RNA-Seq of aging hippocampal microglia.** The tables contain genes selectively increased or decreased in each age group when compared to every other age group combined and genes differentially expressed between 6- and 12-, 18-, or 24-month microglia. The tables contain the average log fold change, the percentage of cells in each group with expression, and the adjusted p-value.

**Table S2 Pseudotime trajectories of aging microglia.** The pseudotime analysis contains the spatial autocorrelation analysis. Moran’s I was used to detect focal expression of genes that were consequently constructed into co-regulated modules using Louvain community analysis. The table contains the adjusted p-value (q-value) and module information.

**Table S3 RNA-Seq analysis of primary microglia treated with LPS and TGFB1 or CX-5461.** The tables contain the RNA-Seq differential expression analysis of control and LPS activated primary microglia, DMSO and CX-5461 treated primary microglia activated by LPS, and DMSO and TGFB1 treated primary microglia activated by LPS, respectively. The tables contain expression values of genes for the samples that were compared along with mean expression values for all samples combined, log fold change between sample groups, and adjusted p-values. in the RNA-Seq differential expression analysis of control and LPS activated primary microglia, DMSO and CX-5461 treated primary microglia activated by LPS, and DMSO and TGFB1 treated primary microglia activated by LPS, respectively.

**Table S4 Differential expression analysis of control and *Tgfb1* cKO hippocampal microglia.** The table contains expression values of genes for the samples that were compared along with mean expression values for all samples combined, log fold change between sample groups, and adjusted p-values.

