## Supplementary figures and images for "Microglia aging in the hippocampus advances through intermediate states that drive activation and cognitive decline"

### Supplementary Figures S1-6

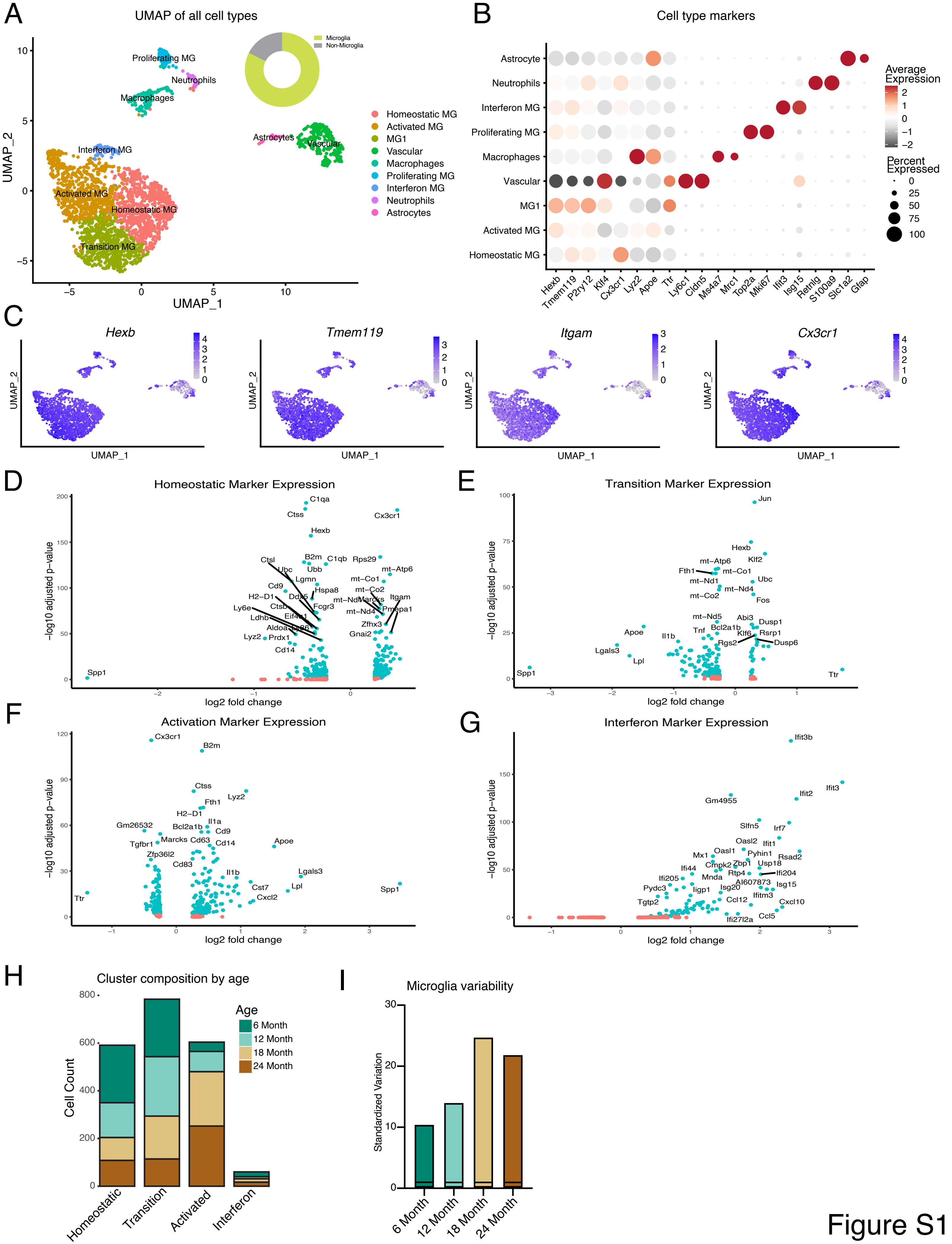

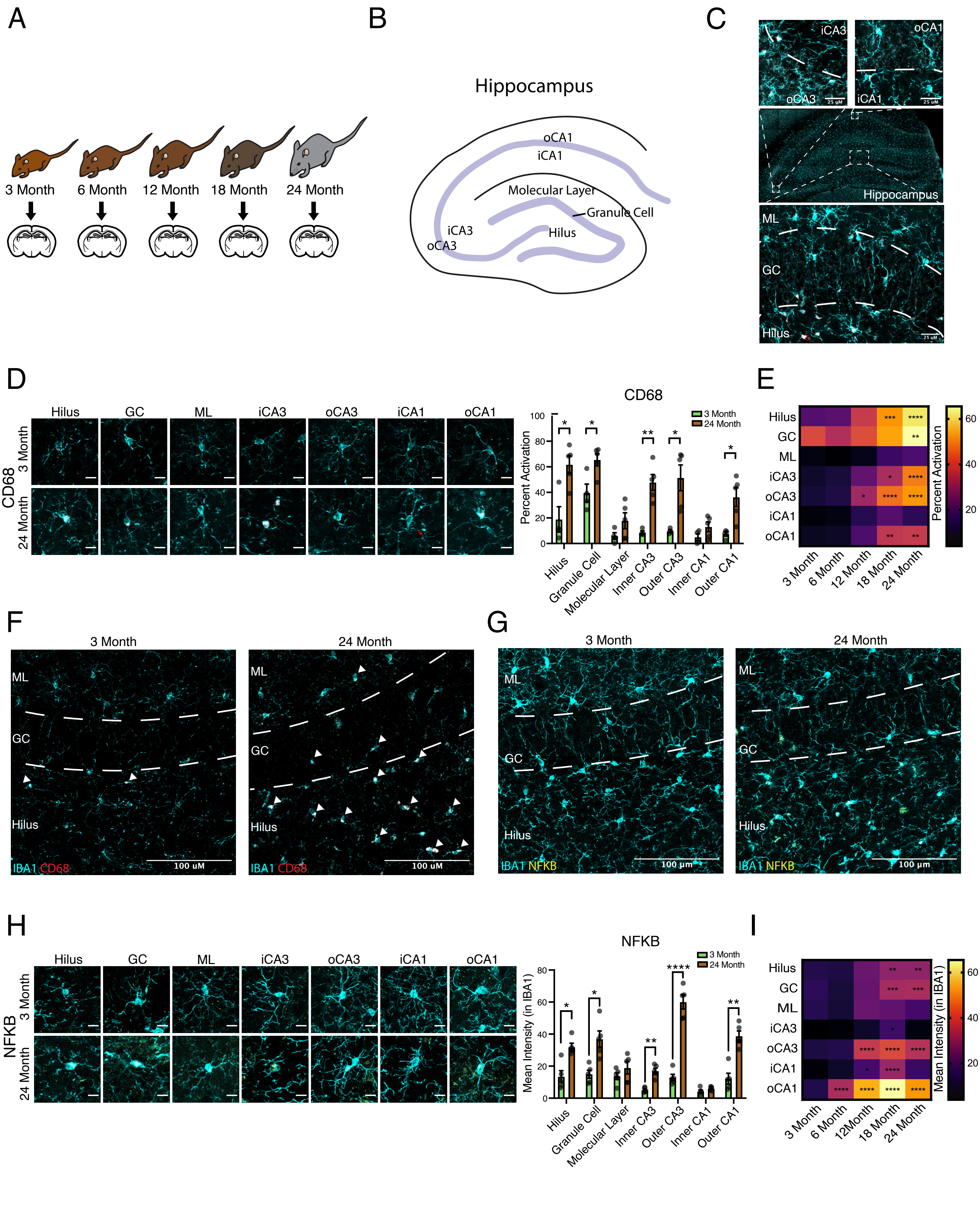

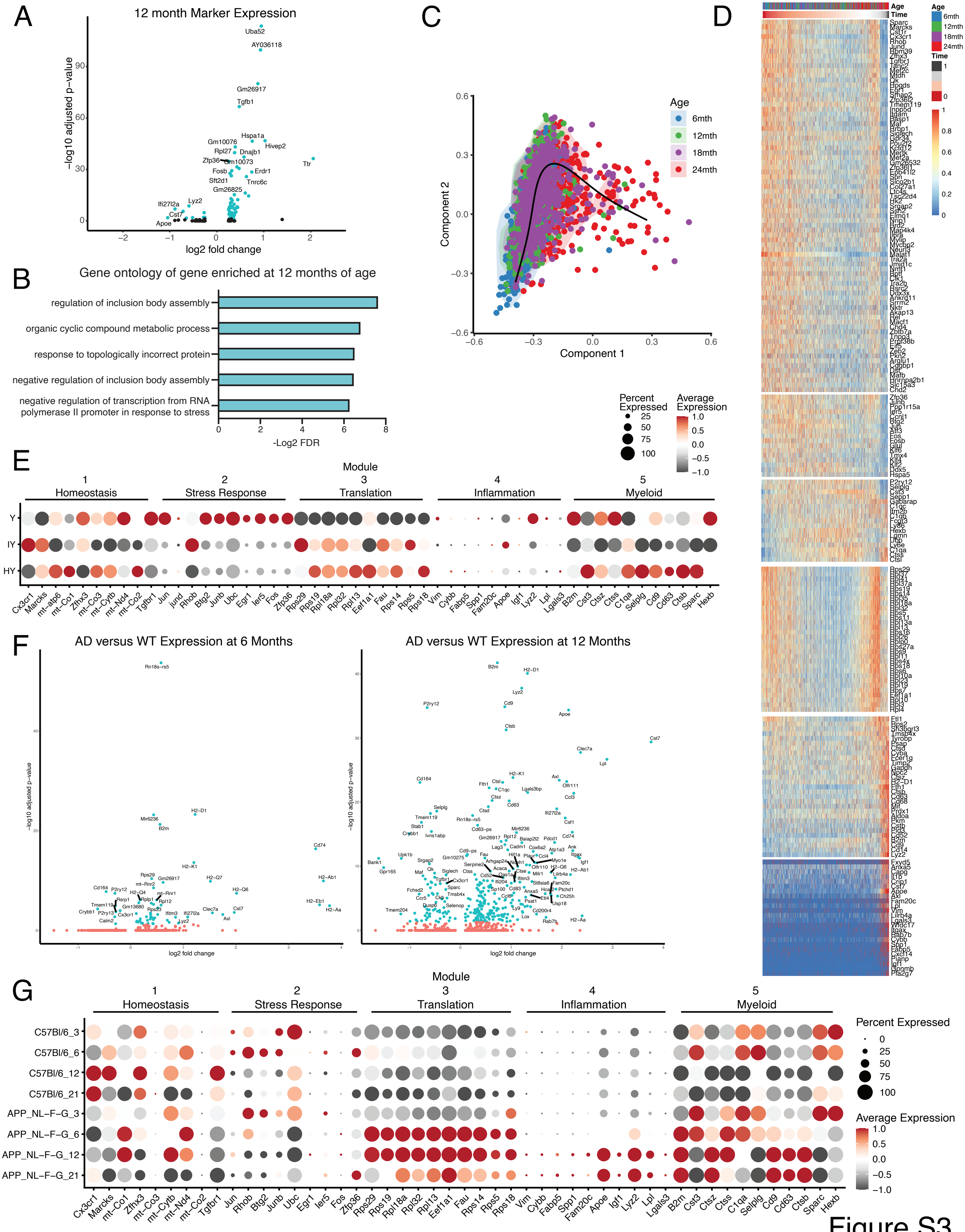

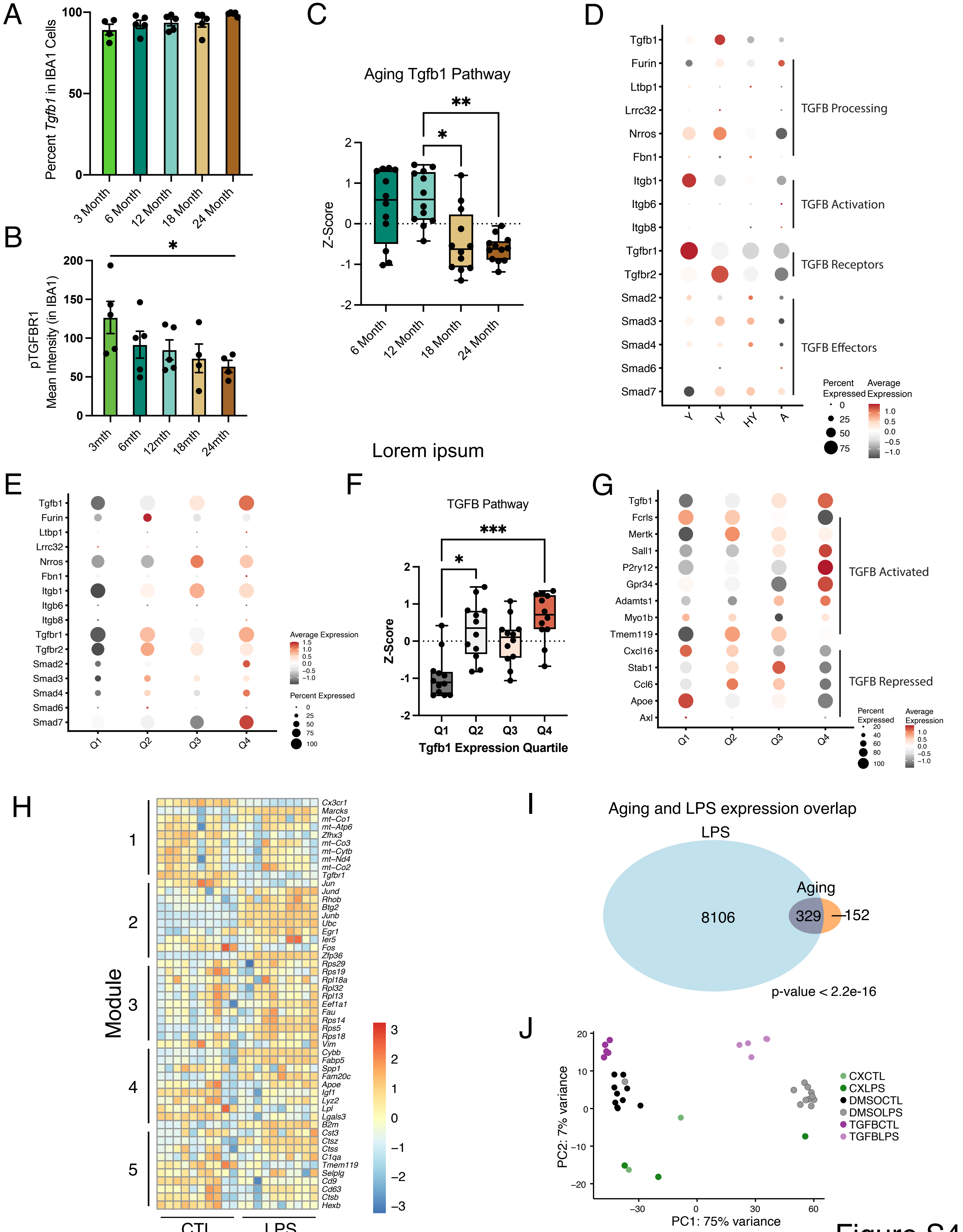

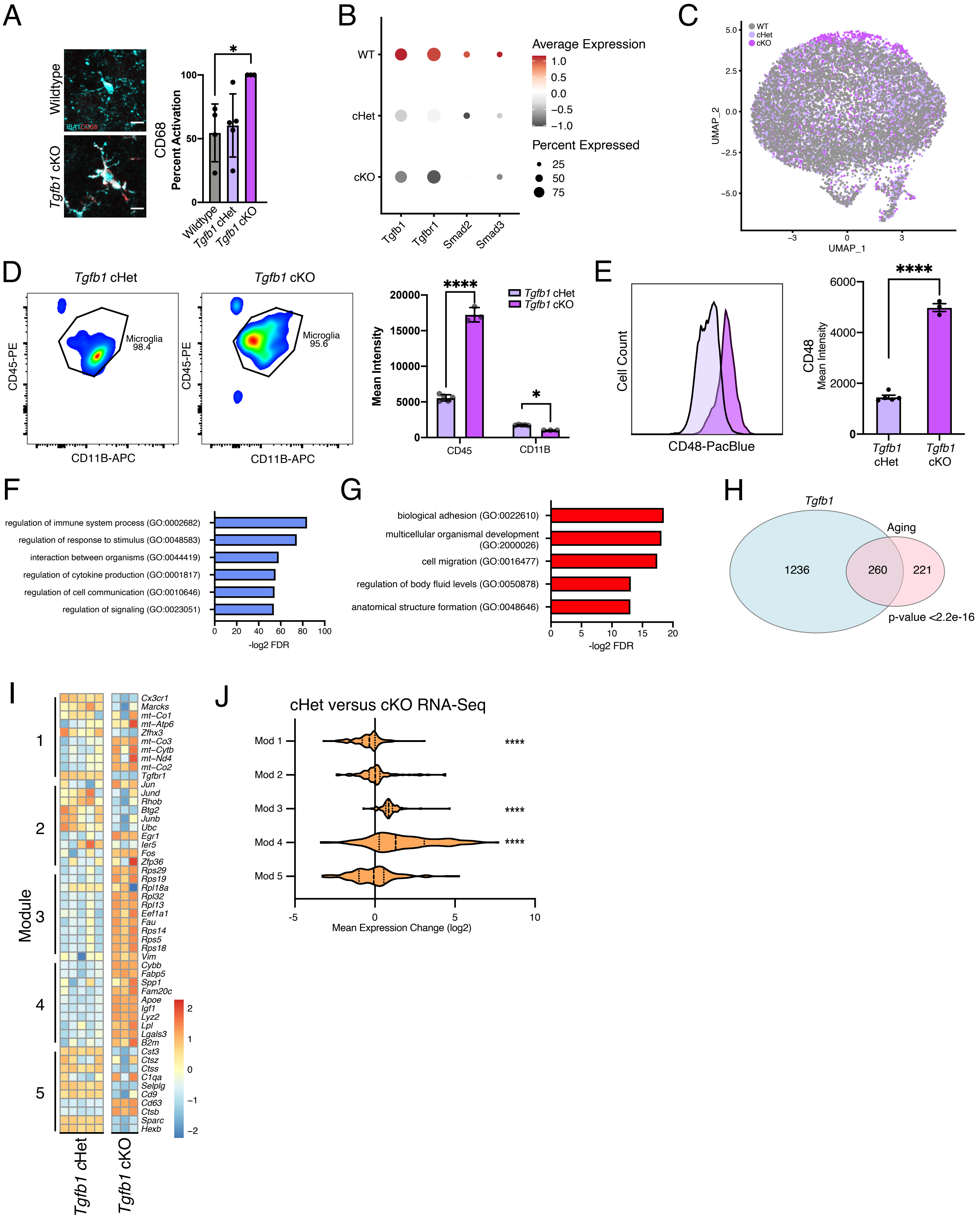

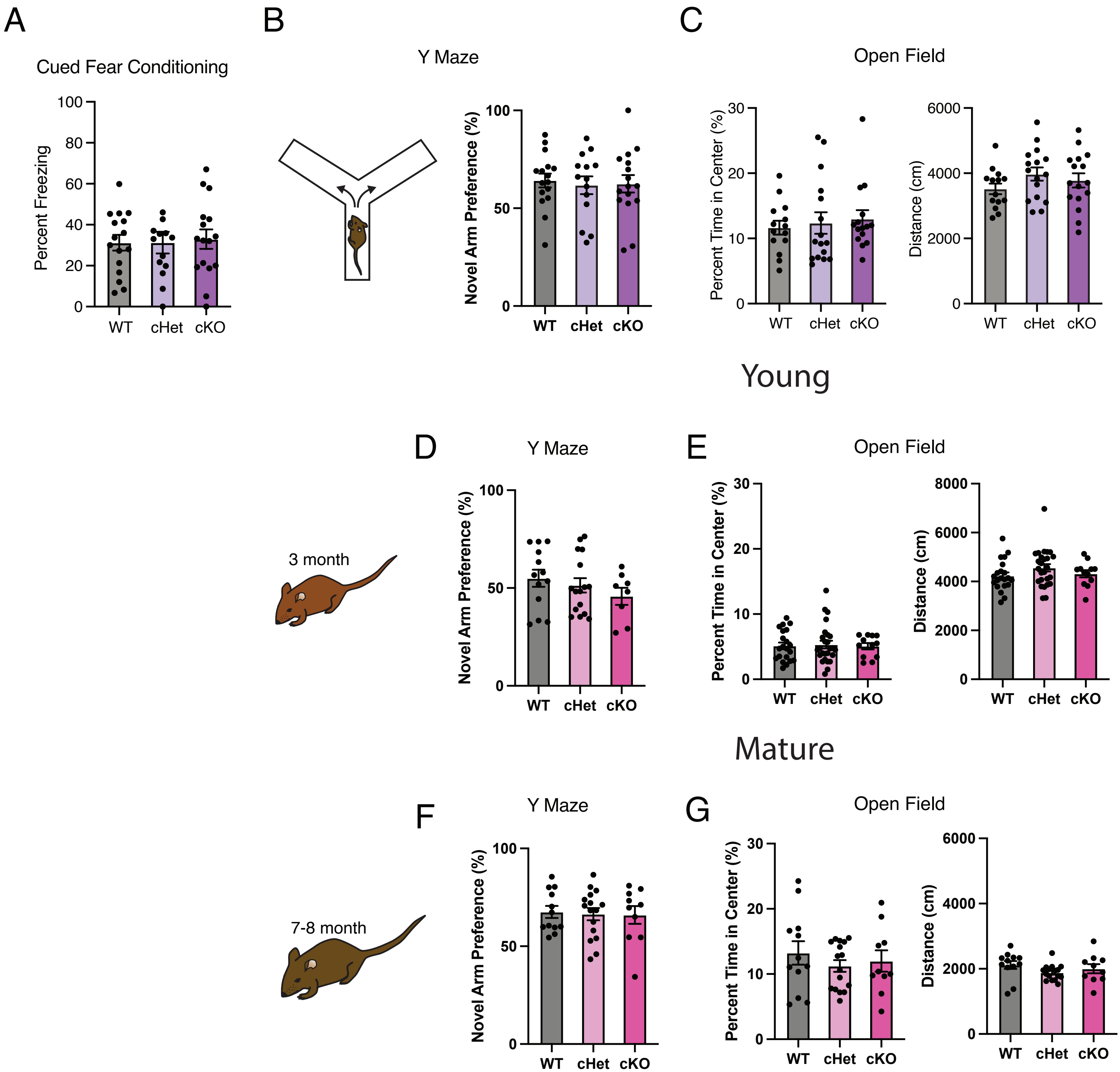
